# REVOLVER: a low-cost automated protein purifier based on parallel preparative gravity column workflows

**DOI:** 10.1101/2021.12.12.472287

**Authors:** Patrick Diep, Jose L. Cadavid, Alexander F. Yakunin, Alison P. McGuigan, Radhakrishnan Mahadevan

## Abstract

Protein purification is a ubiquitous operation in biochemistry and life sciences and represents a key step to producing purified proteins for research (understanding how proteins work) and various applications. The need for scalable and parallel protein purification systems keeps growing due to the increase in throughput in the production of recombinant proteins and in the ever-growing scale of biochemistry research. Therefore, automating the process to handle multiple samples in parallel with minimal human intervention is highly desirable; yet only a handful of such tools have been developed, all of which are closed source and expensive. To address this challenge, we present REVOLVER, a 3D-printed programmable and automatic protein purification system based on gravity-column workflows and controlled by Arduino boards that can be built for under $130 USD. REVOLVER completes a full protein purification process with almost no human intervention and yields results equivalent to those obtained by an experienced biochemist when purifying a real-world protein sample. We further present and describe MULTI-VOLVER, a scalable version of the REVOLVER that allows for parallel purification of up to six samples and can be built for under $250 USD. Both systems will be useful to accelerate protein purification and ultimately link them to bio-foundries for protein characterization and engineering.

**Specifications Table:** 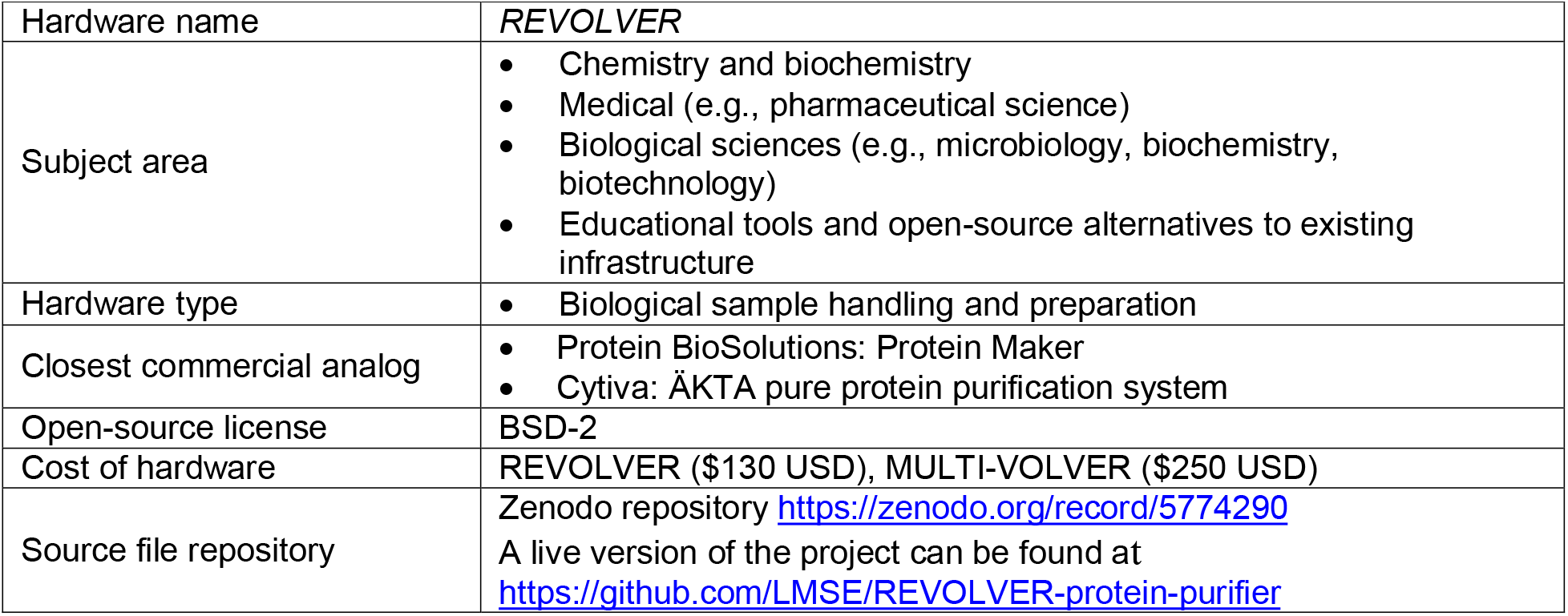

## 1. Hardware in context

To understand living organisms at the molecular level, biomacromolecules are studied in isolation, in cellular systems, and in more complex networks (*e*.*g*., microbial communities or multicellular organisms). Specifically, proteins are one of the major categories of biomacromolecules and their isolation, termed *protein purification*, is a ubiquitous task required in any investigation seeking to learn what a protein looks like and how it works. This is especially true for uncharacterized proteins or those that have been mutated or engineered for some purpose, such as to study diseases or improve enzyme catalysis. After the success of the Human Genome Project and the consequent rise in accessible sequencing technologies by the late 1990s, there existed a plethora of genes encoding for proteins with *no* structural data that would allow scientists to deeply understand the mechanisms by which these proteins functioned [1]. The field of structural genomics (SG) was thus born to address this grand challenge.

From 2000-2015, the field grew immensely with 51 SG facilities solving nearly 15,000 protein structures by X-ray crystallography or nuclear magnetic resonance [1]. To achieve this level of throughput, protein purification needed to become a more industrial endeavor through new technologies and techniques that were adopted during this time, which are still used to this day and will continue to be used in the foreseeable future. There were two features that were needed to purify proteins at scale: parallel workflows, and automation. Prior to SG, proteins were purified using glass gravity columns where a single protein can be purified at a time as described in Fig. 1 [2]. With the need to process more proteins regularly, SG scientists would manually purify proteins in parallel by simultaneously processing several gravity-columns with the appropriate resin packed inside (Fig. 1–5a). There also existed fast protein liquid chromatography systems (FPLC), predominantly from ÄKTA (now owned by Cytiva). To leverage the FPLC’s ability to automatically load the resin with a single protein sample, wash, and elute the target protein (Fig. 1–5b); SG scientists modified these systems (sometimes using existing laboratory equipment like peristaltic pumps) to improve their functionality [3–6]. This type of work is continued to this day by scientists, as well as ÄKTA engineers that have developed newer versions [7–9]. While multiple FPLC systems could be run in parallel, this is an expensive alternative to manually processing several protein purifications by gravity column.

**Fig. 1.**
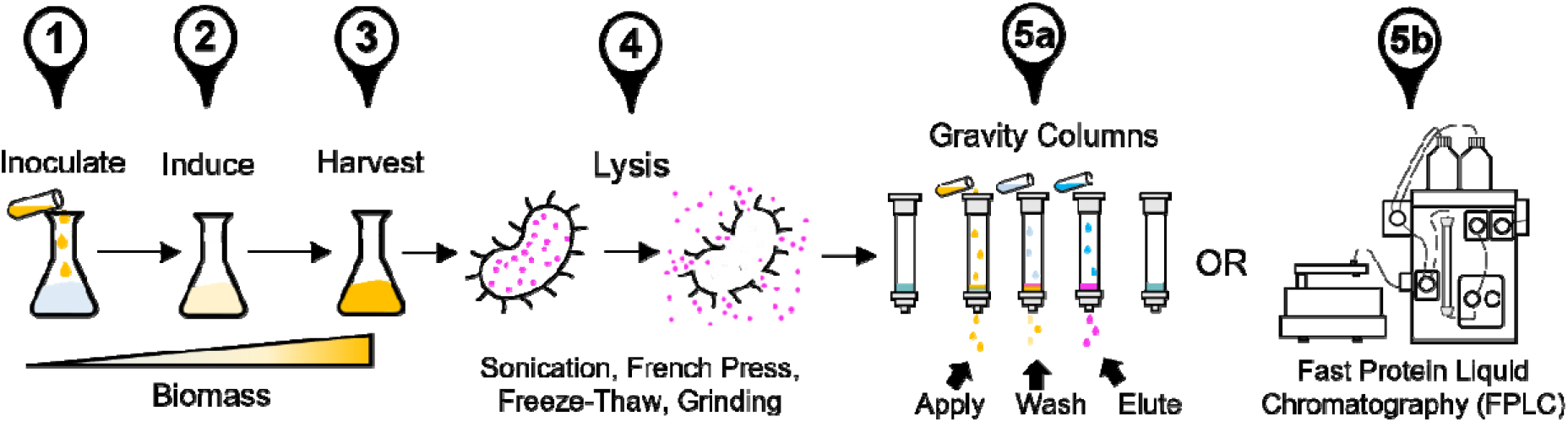
Typical protein purification protocol. We focus on protein purification from *Escherichia coli* modified to express large quantities of a recombinant target protein, highlighted in magenta. Step 1) an overnight culture of the *E. coli* strain is inoculated into a large volume of nutrient media and allowed to grow to medium density (O.D. 0.6-1.0). Step 2) an inducer chemical is added that triggers protein expression. Step 3) continue protein expression by allowing the cell to grow in the presence of the inducer, then harvest the cells by centrifugation. Step 4) use any combination of methods to physically/ chemically lyse the cells to release the target protein. Step 5) remove cellular debris by centrifugation, then apply the supernatant (*i*.*e*., lysate) to either 5a) a gravity column containing the appropriate affinity resin, or 5b) an FPLC system. In (5a), application of the lysate to the resin is followed by a wash to remove bound non-target proteins, then an elution step where changes in some solution condition (*e*.*g*., pH, salt, imidazole) leads to the target protein eluting from the column where it may be collected for further work. The FPLC automatically performs these steps with real-time monitoring of the solution condition and protein detection.

In 2016, the Seattle Structural Genomics Center for Infectious Disease published their work developing the Protein Maker in partnership with automation company Emerald BioSystems (now sold by Protein BioSolutions). The Protein Maker combines both aforementioned features as it can automatically purify up to 24 proteins in parallel [10,11]. The Protein Maker, however, is a commercially supplied solution for automated parallel protein purification that is patent protected (US Patent No. 6818060). Similar technologies include Profinia from Bio-Rad, which is a discontinued product; and the Fluent automated workstation by Tecan, which is extremely expensive. While both gravity columns and FPLC are used to this day, to our knowledge there does not exist an inexpensive open-source solution that achieves automatic and parallel protein purification. Therefore, we sought to design, build, and test a scalable system that can perform the basic functions of a paralleled FPLC system (*i*.*e*., the Protein Maker). Specifically, we want to process cell lysates by applying them to an affinity resin, washing the resin, and eluting the protein from the resin (Fig. 1–5a). We call our inventions the REVOLVER and the MULTI-VOLVER.

## 2. Hardware description

In this section we first describe the design of REVOLVER, which is comprised of several components that can be independently 3D-printed and fitted with the firmware to form one functional device (Fig. 2, Section 2.1). Several REVOLVERs can be connected with additional components and firmware for automated parallel protein purification. We call this combination the MULTI-VOLVER, which we discuss in a subsequent section (Fig. 3, Section 2.3). Our printed circuit board (PCB) design is discussed in Section 2.3; and the communication protocol between the Arduino Nano microcontrollers and their interaction with the user is discussed in Section 2.4. While we do our best to explain various concepts underlying these two inventions herein, we strongly suggest the reader first consider watching Supp.Video 1 – 4 to see the systems in action.

**Fig. 2.**
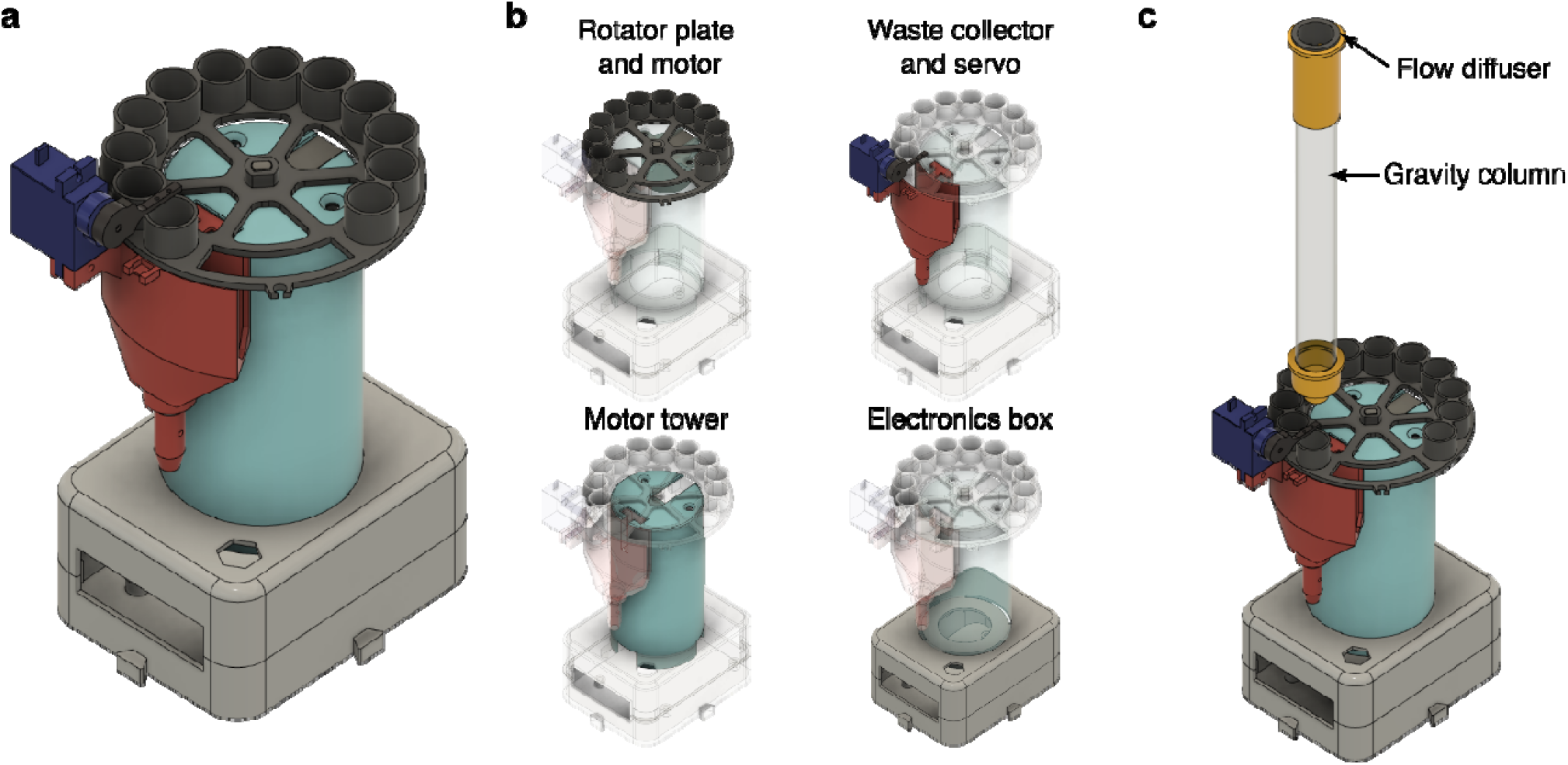
Design of the REVOLVER protein purification system. (a) Concept, (b) Overview of the bodies comprising one REVOLVER device, (c) Schematic of the REVOLVER system with a gravity column and a flow diffuser.

**Fig. 3.**
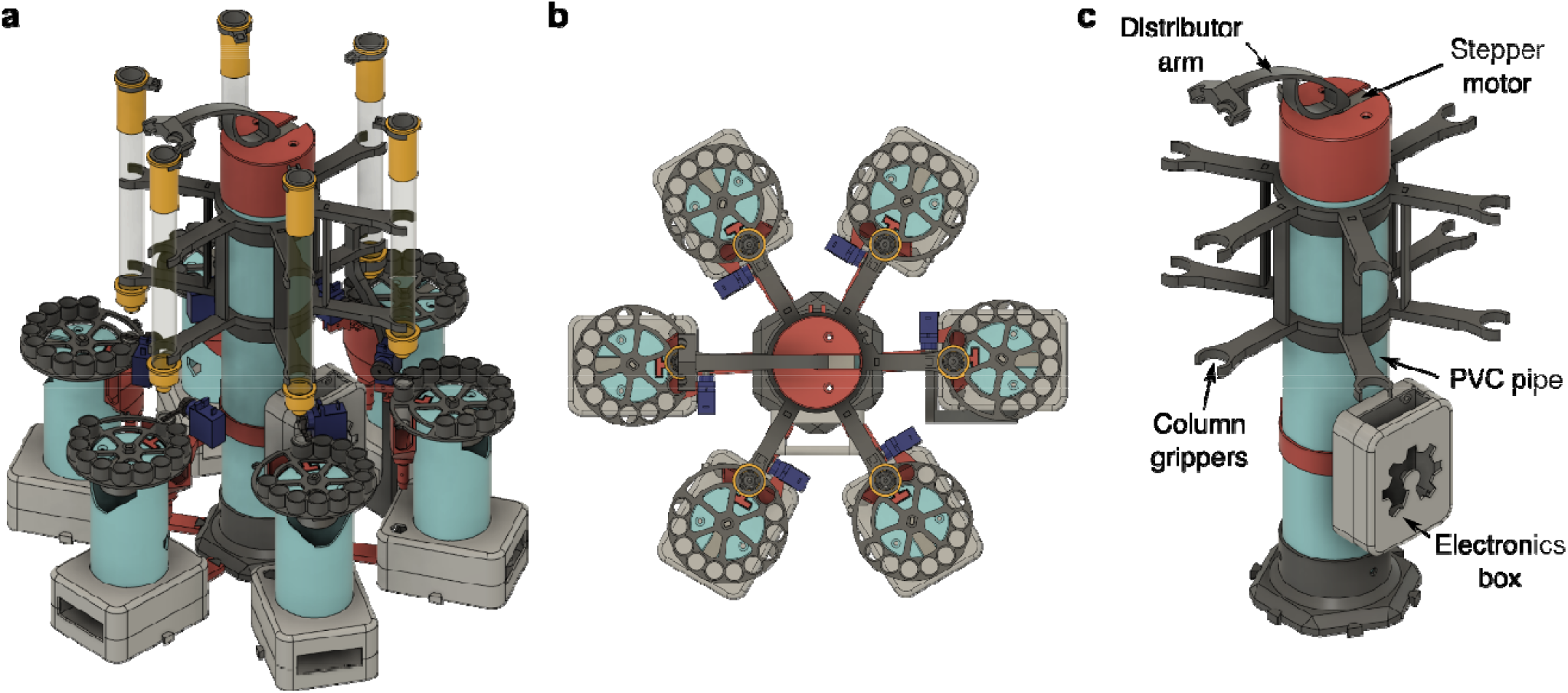
Design of the MULTI-VOLVER. (a) Concept of MULTI-VOLVER with six REVOLVERs, (b) Top-down view, (c) Components specific to the distributor in the MULTI-VOLVER configuration. The 3D-printed column grippers allow for precise positioning of the individual gravity columns. Note the electronics box here is different from the REVOLVER version.

**Figure 4.**
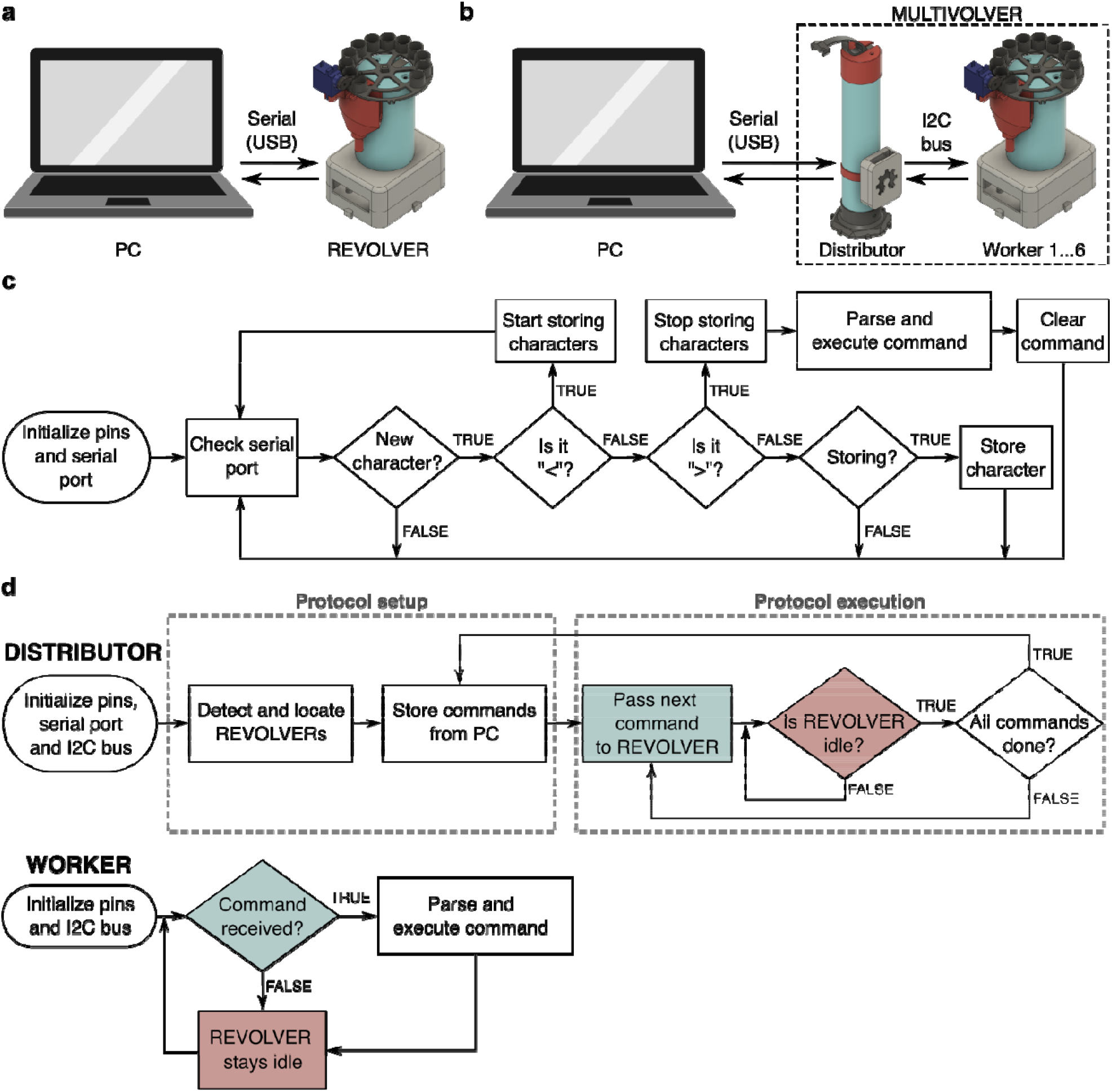
REVOLVER and MULTI-VOLVER Communication Protocol. (a) The user interface with the REVOLVER is through a serial USB connection with a personal computer (PC), (b) The user interface with the MULTI-VOLVER is through a serial USB connection a PC and the Distributor, which is separately connected to the individuals REVOLVERs (*viz*., the Workers), (c) Algorithm for storing and parsing commands in the REVOLVER, (d) Algorithm for the MULTI-VOLVER’s Distributor and individual REVOLVERs (*viz*., the Workers). Commands are stored with an algorithm similar to that in panel (c). Blocks with the same color in the Distributor and Workers denote communicatio between the devices via I2C

**Figure 5.**
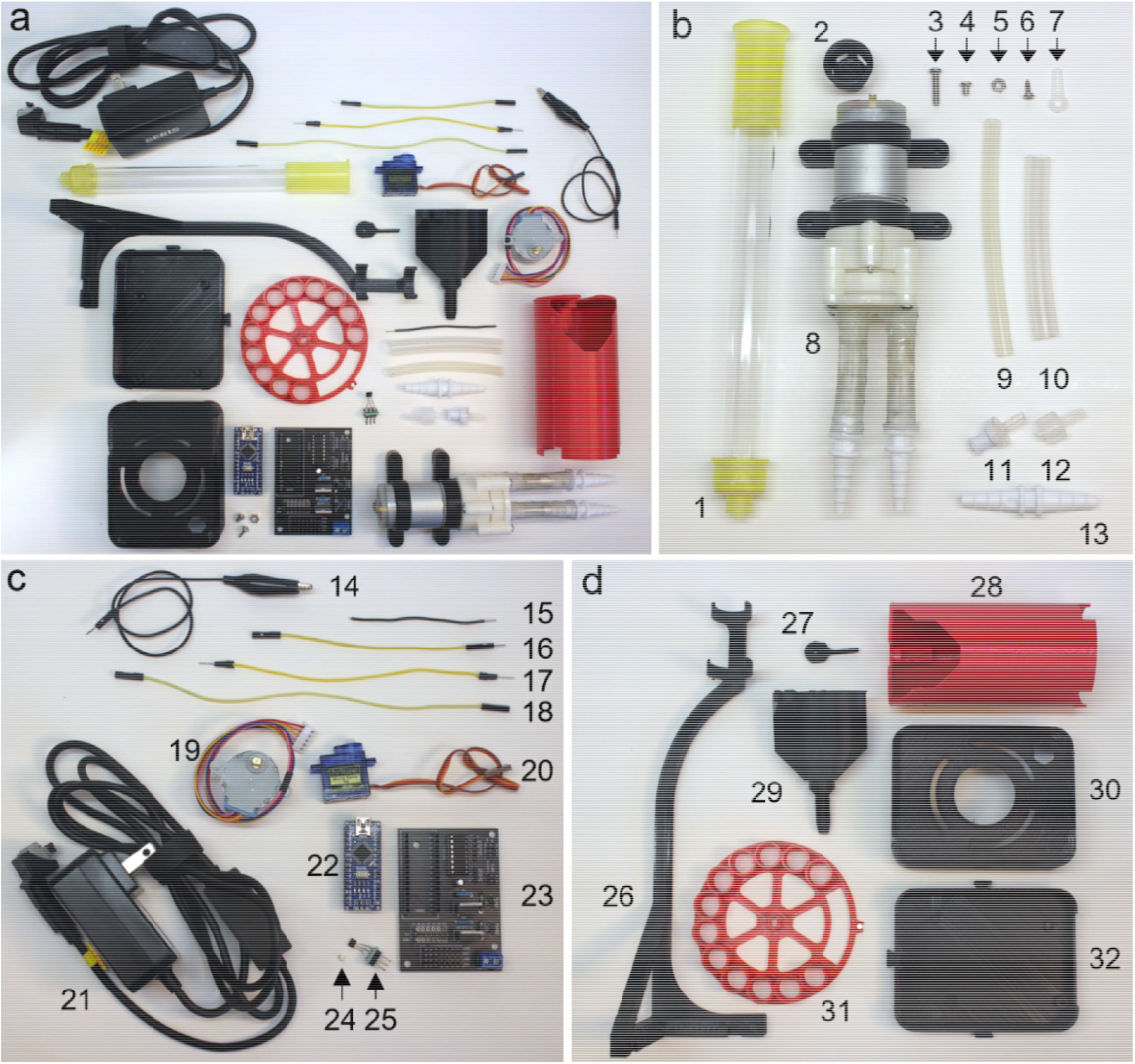
Top-down view of unique parts for the REVOLVER. (**a**) Overview of all unique parts, (**b**) Parts related t liquids: 1) column, 2) a8 – flow distributor, 3) 12 mm M3 screw, 4) 6 mm M3 screw, 5) M3 nut, 6) 6 mm Phillips screw, 7) servo horn, 8) DC pumps with fittings for small tubing (*cf*. Supp.Fig. 1), 9) small tubing, 10) large tubing, 11) 1-wa valve, 12) valve adaptor, 13) tube connector; (**c**) Electronics components: 14) alligators (M/F), 15) hookup wire, 16) jumper (MF), 17) jumper (MM), 18) jumper (FF), 19) stepper, 20) servo, 21) power supply, 22) nano, 23) PCB (soldered), 24) magnet, 25) Hall sensor, and (**d**) 3D-printed components: 26) a9 – column grip, 27) a5 – servo arm, 28) a1 – tower, 29) a3 – waste collector, 30) a6 – box top, 31) a2 – plate, 32) a7 – box bottom

**Figure 6.**
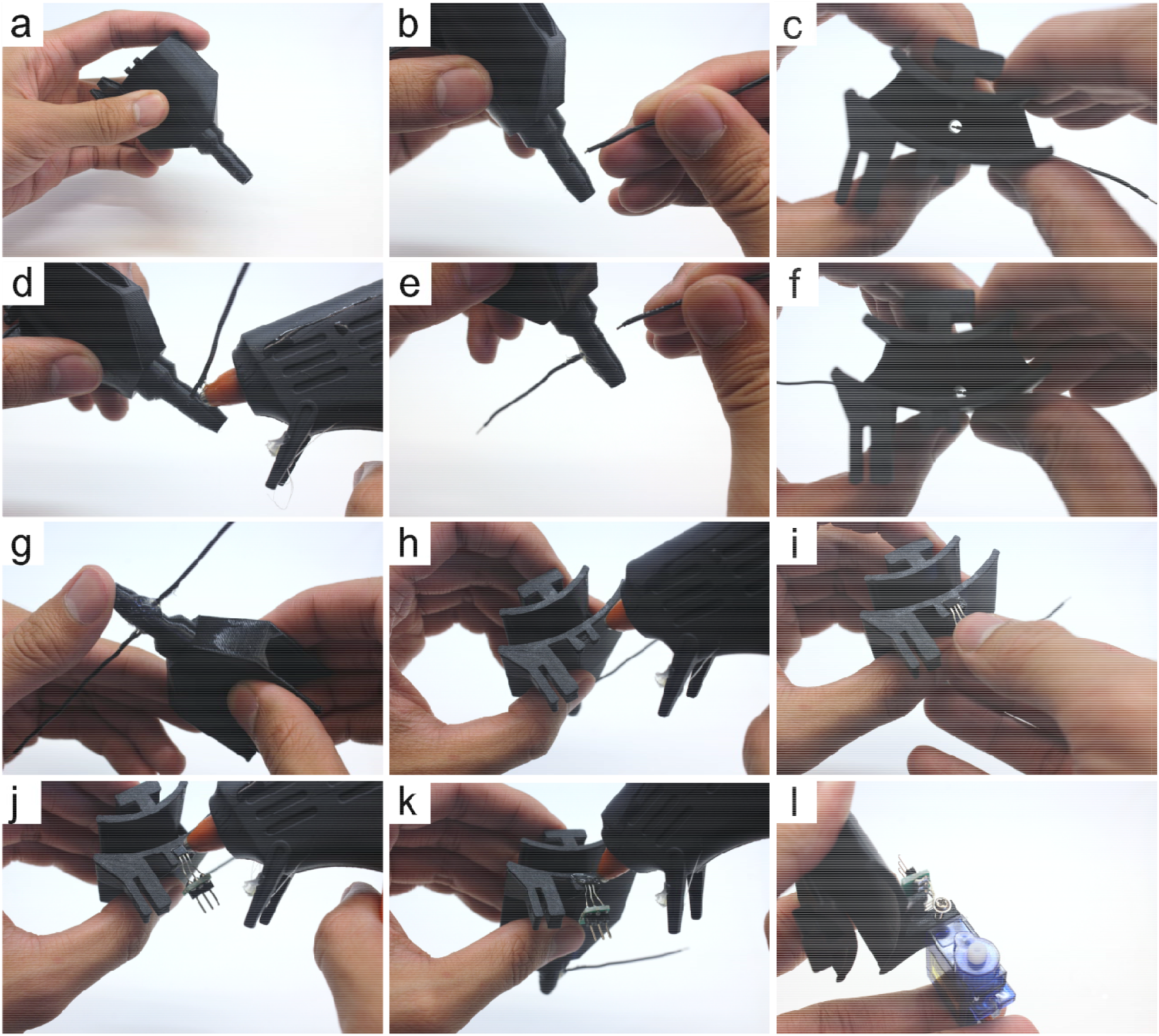
Step-by-step guide to assemble the waste collector. Step (a): remove any filament stringing inside the waste collector (Part 29) Step (b): cut 8 cm of hookup wire (Part 15), strip 1.5 mm and 8 mm of the insulation on opposite ends, then insert the 1.5 mm stripped end into the holes on the side of the waste collector Step (c): push the wire in until roughly 1 mm of the insulation is inside. The exposed wire should not be touching the wall of the waste collector. Step (d): apply hot-glue to create a seal between the hook-up wire and the waste collector Step (e): repeat steps (b) and (c) for the other hole Step (f): check to make sure that both 1.5 mm stripped ends inserted into the waste collector do not touch each other or the walls, and that they both have roughly 1 mm of insulation inside. This ensures that the waste sensor is triggered only when the column is actively dripping solution into the waste collector Step (g): repeat step (d) to create a seal for the other hole Step (h): apply hot-glue to the short slot for the Hall sensor Step (i): insert the Hall sensor (Part 25) into the short slot. Take note of which side you put face down as this becomes relevant in Section 5.1.4. Steps (d) and (e) Step (j): apply hot-glue on top of the Hall sensor Step (k): use the tip of the hot-glue gun to create a seal around the Hall sensor and its metal pins. Thi ensures liquid does not interfere with the Hall sensor’s function. Step (l): insert the servo into the long slot such that the motor axle is on the top, then screw it in place using Part 6.

### 2.1. REVOLVER: single sample fraction collector

The objective of the REVOLVER (Fig. 2a) is to take the user’s cell lysate as input and produce small aliquots of purified protein in microfuge tubes as outputs with no human intervention. To do this, the REVOLVER must move various solutions through the resin packed into the gravity-column (herein termed ‘column’ for brevity). Specifically, after manual application of the protein sample to the resin (*viz*., users must first pour the cell lysate into the gravity-column), REVOLVER must detect when the cell lysate has finished flowing out of the column, then automatically wash the protein-bound resin and elute the target protein into a set of microfuge tubes for downstream processing. REVOLVER’s current hardware is designed specifically for these core steps using the following features: 1) flushed joints requiring minimal screws for simple assembly, 2) closed-loop functionality for automatic handling of solutions, and 3) flow reduction to limit disturbances in the packing of the resin.

#### 2.1.1. Simplified assembly

First, REVOLVER’s bodies (Fig. 2b) are designed to fit into each other using joints (*e*.*g*., tongue-and-groove, double-lap butt). The waste collector fitted with the servo that holds the tube sensor is first combined with the motor tower, which is together inserted into the slits available on the top of the electronics box. If desired, hot glue is sufficient to lock these bodies together (*e*.*g*., the motor tower and electronics box). Screws are only needed to attach the motors to the bodies since these electronics make precise movements for which REVOLVER’s functionality depends on. An optional Hall sensor can be incorporated into the waste collector to auto-home (*i*.*e*., centre) the rotator plate. The plate is inserted onto the motor tower’s stepper. While not shown in Fig. 2, the gravity column can be held using a single-device column gripper that we have designed if the user does not have a metal rod-plate apparatus. These design features were intentional for the purpose of minimizing the number of parts and assembly actions. Detailed assembly instructions can be found in Section 5.0 (Fig. 7–14, Supp.Fig. 1-4).

**Figure 7.**
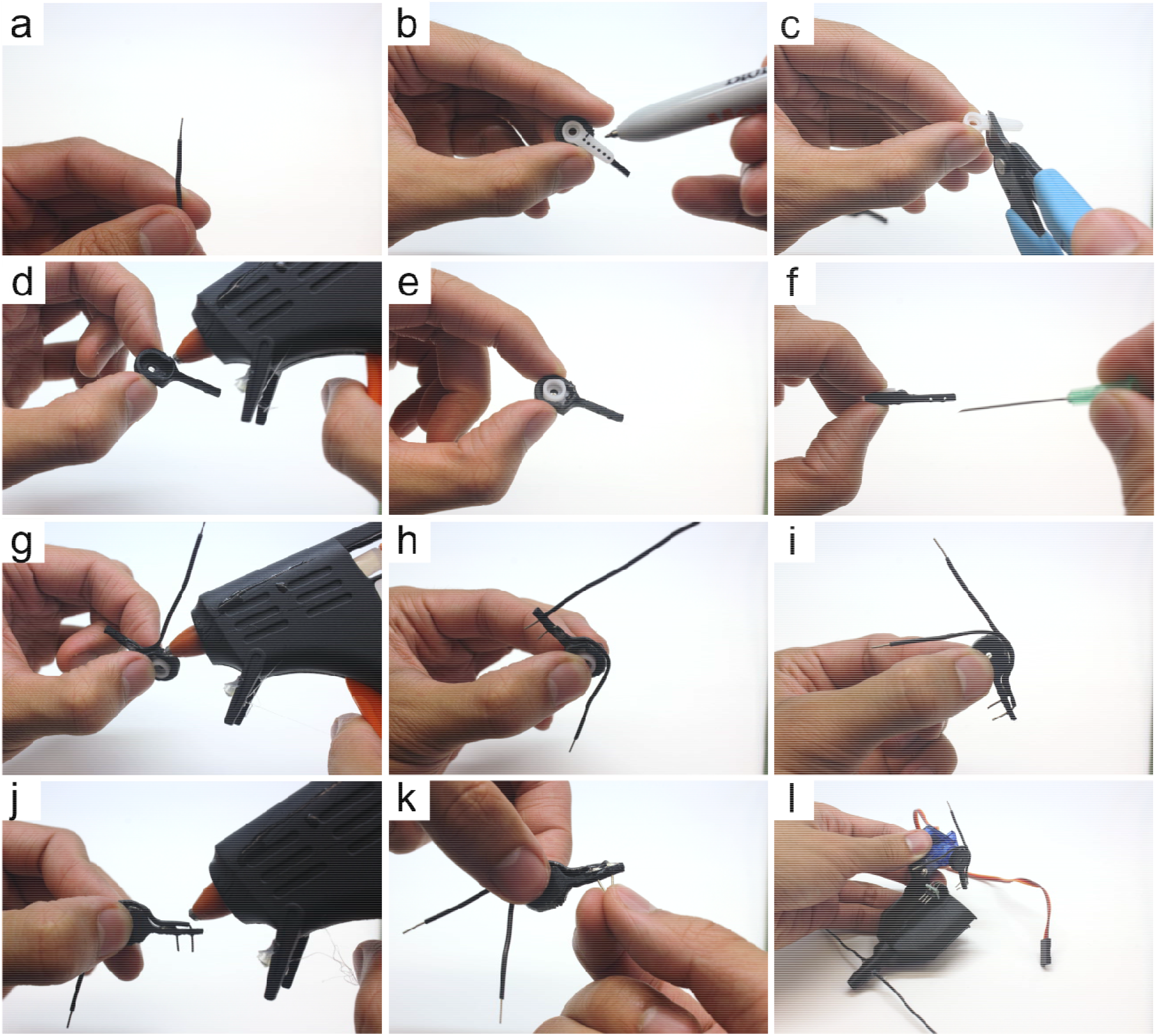
Step-by-step guide to assemble the tube sensor. Step (a): cut two 8-10 cm of hookup wire, strip 8 mm of the insulation on both ends for both pieces Step (b): align the horn (Part 7) with the servo arm (Part 27). Draw a line to indicate where to make a cut to be able to press the horn into the servo arm’s slot Step (c): cut the horn Step (d): apply hot-glue in the servo arm’s slot Step (e): insert the cut horn in the servo arm’s slot Step (f): this step is optional, but depending on the user’s 3D-printer, the holes in the servo arm ma print too small as it did for ours. We recommend using a 21G conventional syringe needles (*e*.*g*., BD ID: 305165) to *carefully* press through the original holes to widen them Step (g): insert one 8mm stripped end (from Step (a)) into the hole closest to the servo arm’s slot. Appl hot-glue and pressure the wire down along the curvature of the servo arm Step (h): insert the other 8 mm stripped end of the other wire (from Step (a)) into the distal hole. Appl hot-glue and press the wire down along the curvature of the servo arm. The two wires should be running along the curvature in a parallel fashion. Step (i): use hot-glue to secure them in place Step (j): apply additional hot-glue where the wires enter the holes of the servo arm to create a seal that prevents liquids from electrically connecting the two pins Step (k): press the pins together, leaving a 2-3 mm gap at their ends Step (l): attach the completed tube sensor to the servo from Section 5.1.1 Step (l) such that the wires point inwardly into the waste collector

**Figure 8.**
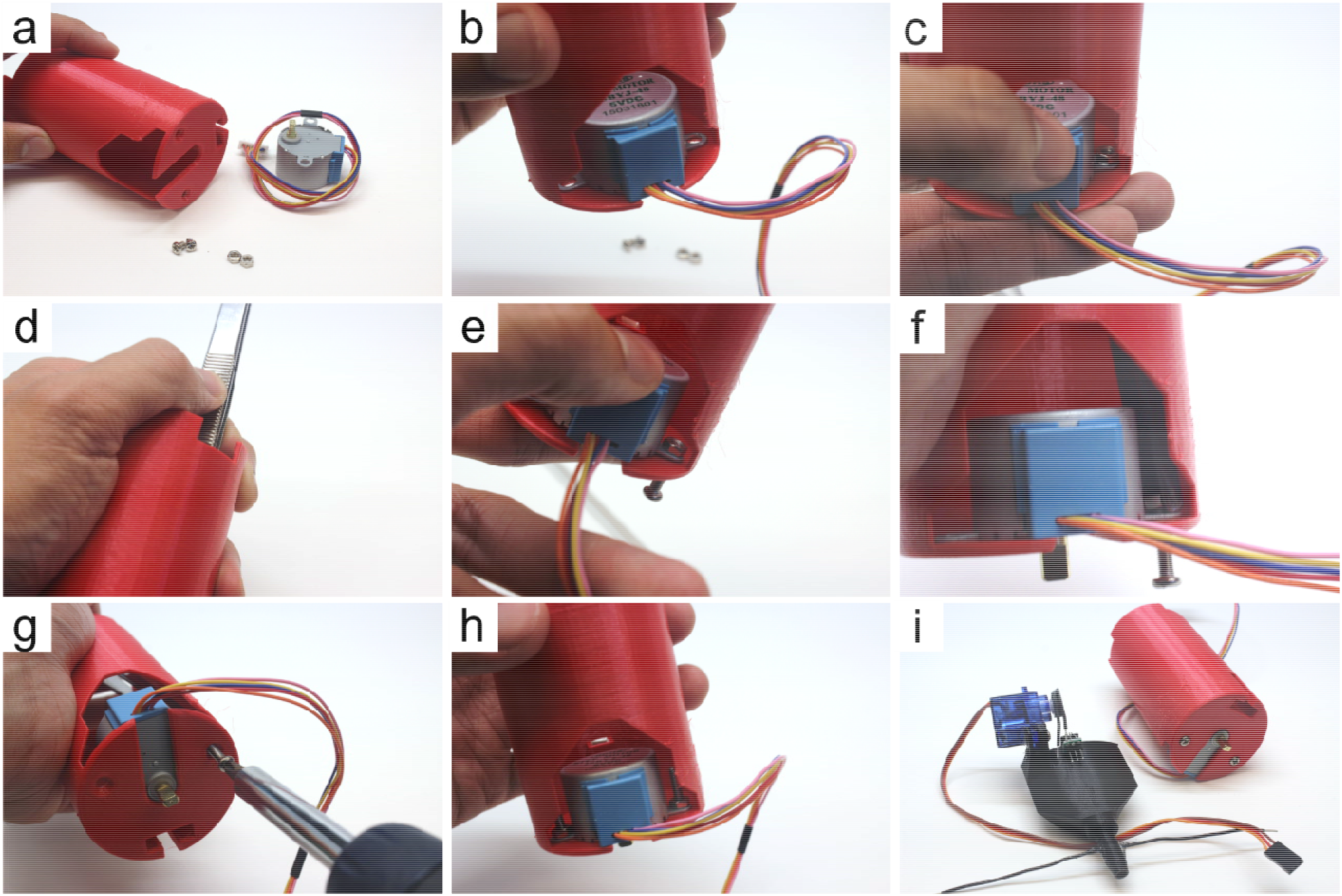
Step-by-step guide to assemble the motor tower. Step (a): remove any stringing inside the tower (Part 28) Step (b): place stepper (Part 19) into the slot on the top of the tower and align the holes of the stepper with the holes in the tower Step (c): place an M3 nut (Part 5) on top of the stepper hole in alignment with the tower’s holes Step (d): using long tweezers, hold the M3 nut in place with one hand, maintaining the alignment with the stepper and tower holes. We recommend using tape (or hot-glue) to help hold the stepper in place temporarily Step (e): screw in the 12 mm M3 screw (Part 3) by hand enough so that it grips the M3 nut Step (f): use the tweezers again to hold the nut in position Step (g): use a screwdriver to tight the screw and the nut to lock the stepper in place with the tower Step (h): repeat steps (c-g) for the other hole Step (i): place the assembled waste collector fitted with the tube sensor aside with the motor tower

**Figure 9.**
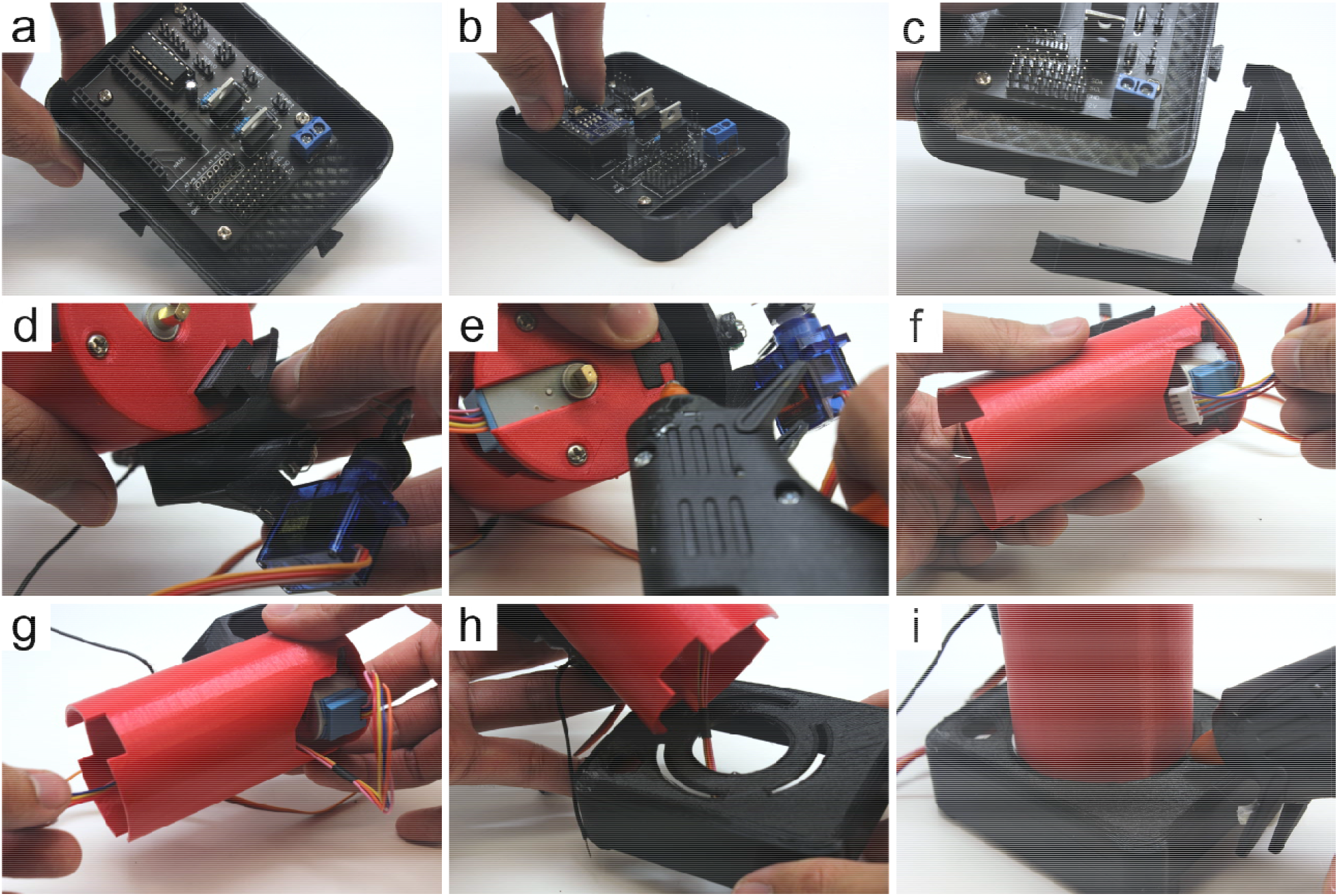
Step-by-step guide to assemble the major components of the REVOLVER. Step (a): attach the soldered PCB (Part 23) to the bottom portion of the electronics box (Part 32) using 6 mm M3 screws (Part 4) Step (b): insert the nano (Part 22) into the soldered PCB Step (c): attach the column grip (Part 26) to the latches outside of the bottom portion of the electronics box (Part 32) Step (d): combine the assembled waste collector from Section 5.1.1 / 5.1.2. with the motor tower from Section 5.1.3. using the joint of the waste collector Step (e): this step is optional, but the user’s 3D-printer tolerances may differ from ours. We recommend applying hot-glue to the connection formed in step (d) to fully lock the two components together Step (f): tuck the stepper’s wiring into the motor tower Step (g): pull the stepper’s wiring through the motor tower Step (h): and pull the stepper’s wiring through the circular hole of the electronics box (Part 30) Step (i): insert the grooves of the motor tower into the top of the electronics box and apply hot-glue to lock the components together

**Figure 10.**
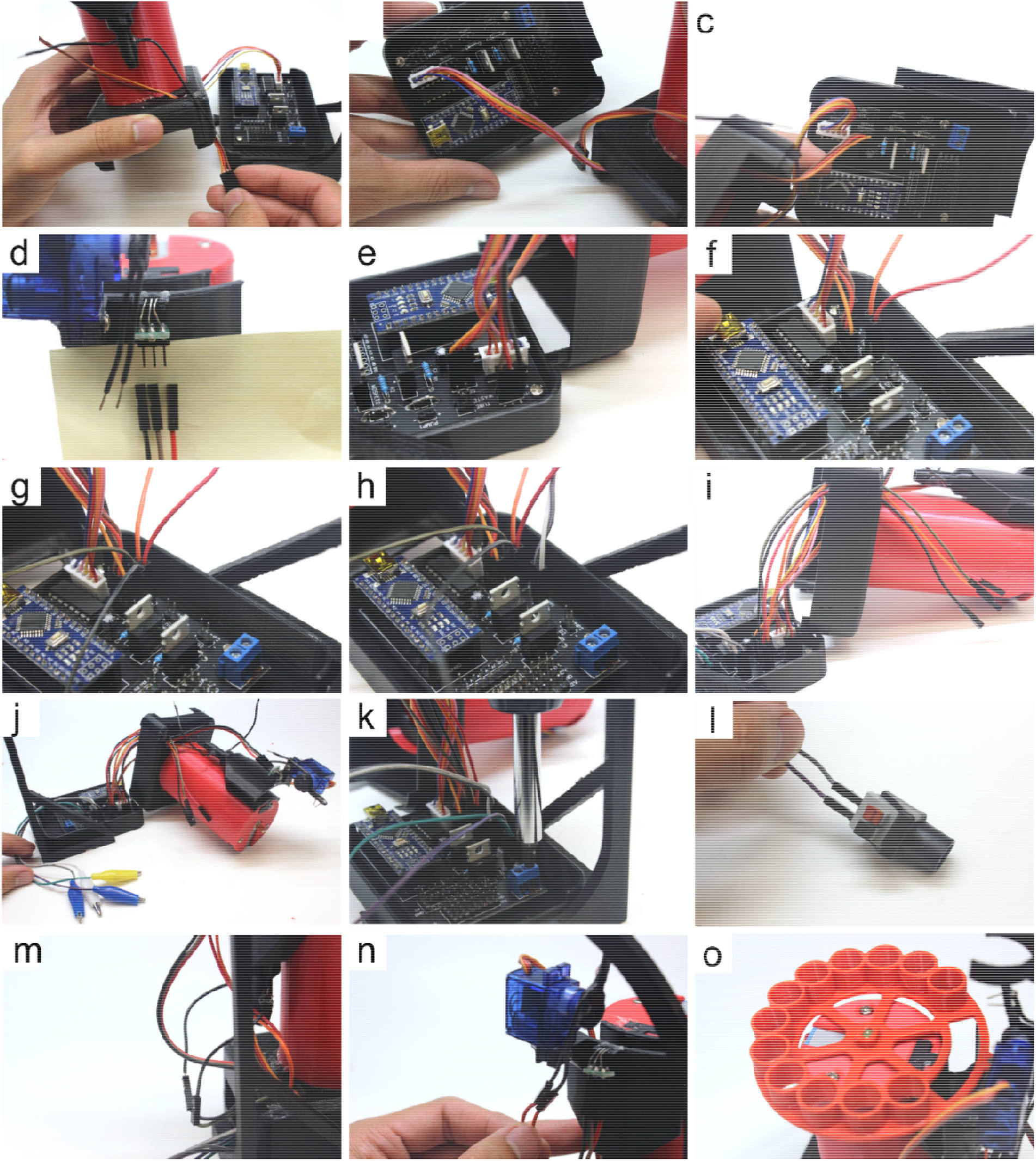
Step-by-step guide to combine the hardware with the firmware. Step (i): pull the unattached jumper (FF) wires from steps (f) and (g) through the small hole of the electronics box Step (a): pull the alligators to one end of the electronics box and check that all remaining wires (attached on unattached at this point) are pulled through the electronics box’s small hole. Attach each pair of alligators to a pump’s positive and negative clips (Part 8) Step (j): loosen the screw block of the PCB and insert one jumper (MM) wire (Part 17) into each positive and negative terminal, then re-tighten the screw block to lock the wires in place Step (k): connect the ends of the jumper (MM) wires from the previous step to the power supply’s adaptor piece, then connect the adaptor to the power supply (Part 21) Step (l): connect the jumper (FF) wires from step (f) to the tube sensor’s exposed 8 mm stripped ends from Section 5.1.2. step (l). The polarity here will not affect the function Step (m): connect the jumper (FF) wires from step (g) to the waste sensor’s exposed 8 mm stripped ends from Section 5.1.1. step (g). The polarity here will not affect the function Step (n): place the plate (Part 31) onto the servo (Part 19) of the motor tower. Ensure that a magnet (Part 24) has been placed into its slot in the plate.

**Figure 11.**
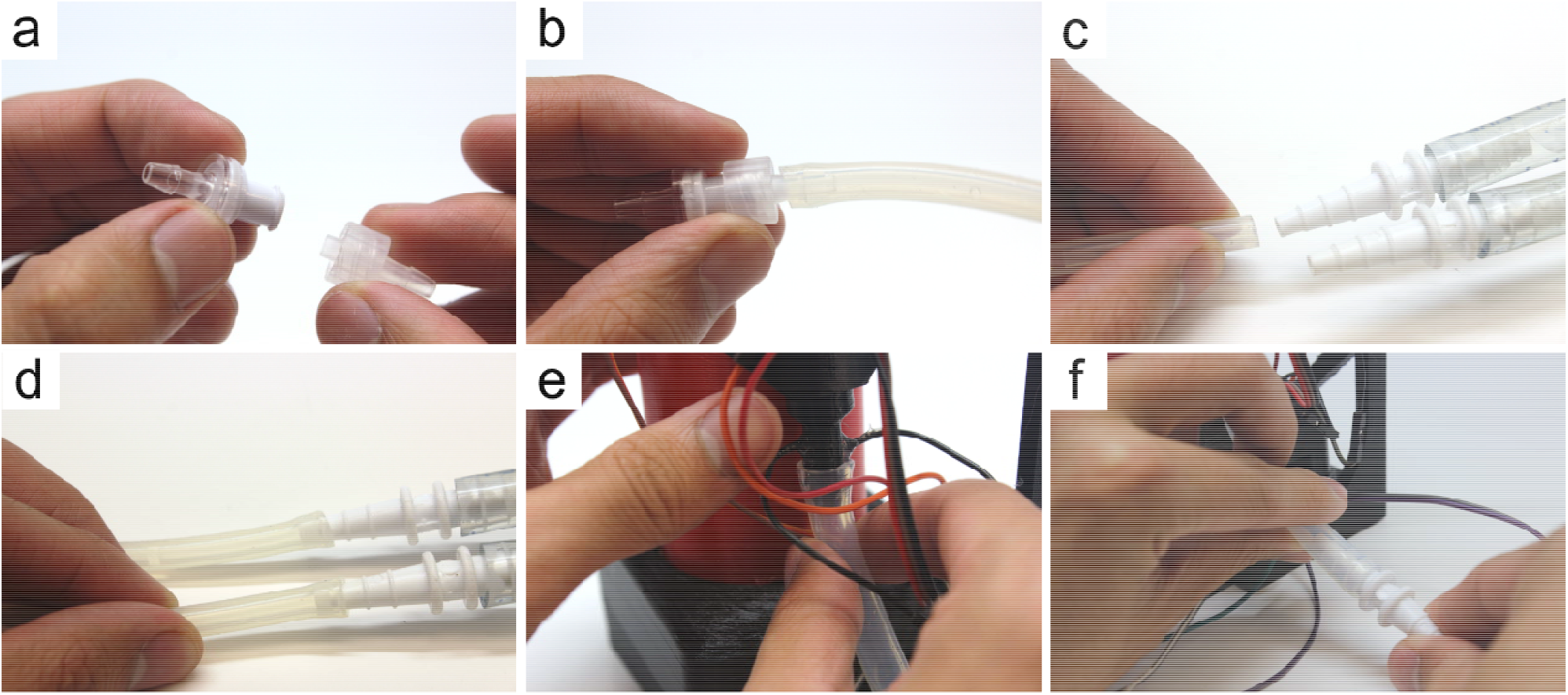
Step-by-step guide to wire the hardware with the firmware. In this section, each pump is dedicated to transferring solutions from one container to the column. Two pumps exist since the protein purification protocol requires a wash and elution solution. Step (a): connect the 1-way valve (Part 11) with the valve adaptor (Part 12) Step (b): insert the valve adaptor from the previous step into a small tubing piece (Part 9) cut to the user’ desired length (recommended 45-60 cm) Step (c): insert the tube connector of the pump (Part 8) where fluids are pushed out (positive pressure end) to the end of the tubing piece from the previous step Step (d): cut another tubing piece from Part 9 to a desired length (recommended 45-60 cm) and insert the negative pressure end of the pump to it. The end of this tubing piece is placed into the wash solution. Repeat steps (a-d) for the other pump to be used for the elution solution Step (e): cut 10 cm of the large tubing (Part 10), and connect one end to the bottom of the waste collector Step (f): attach a tube connector to the other end of the large tubing piece from the previous step. An additional small tubing piece (cut to approximately 45-60 cm) can be attached to the other end of the tube connector, which can be drained into a waste receptacle

**Figure 12.**
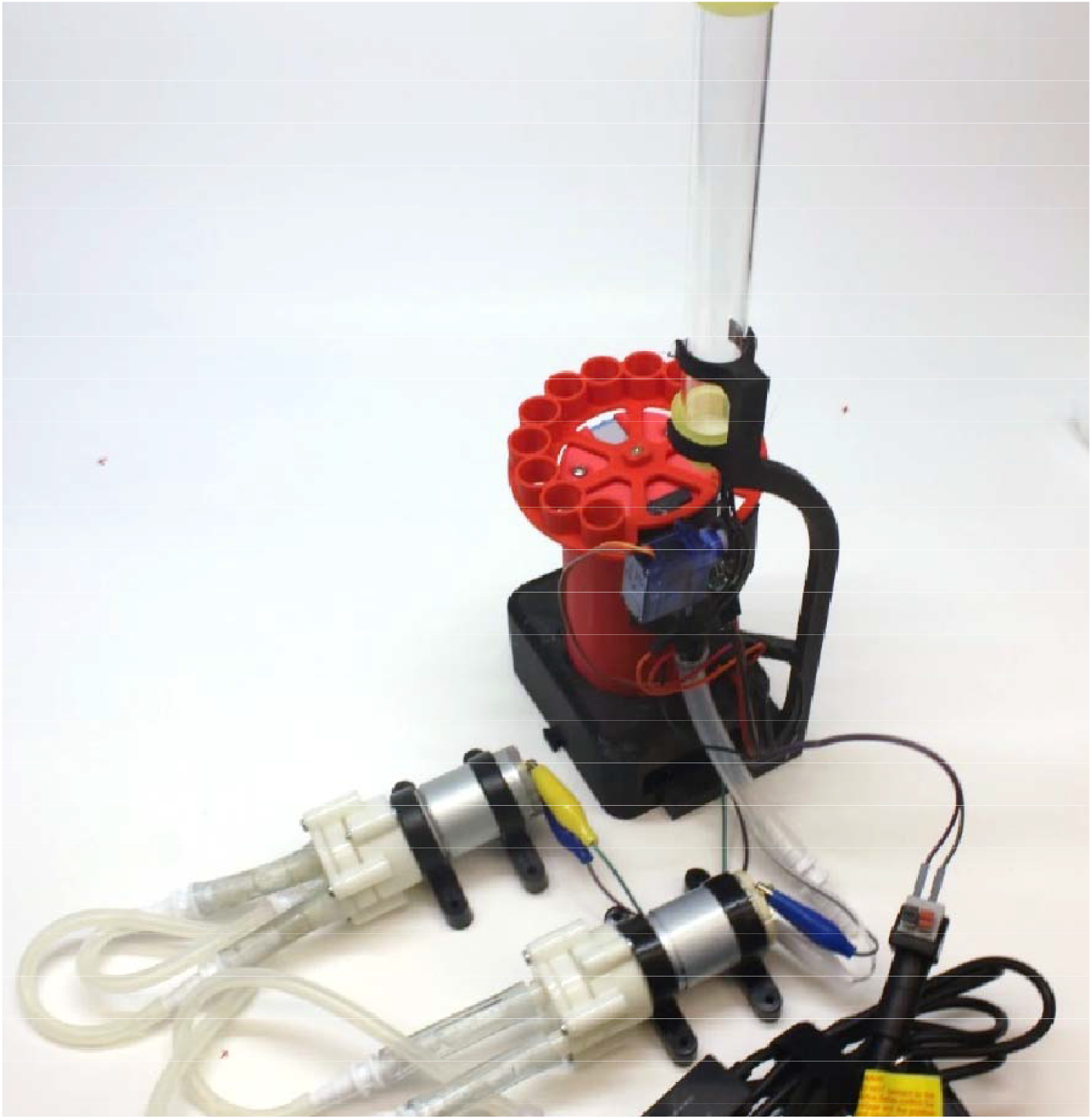
Fully-assembled REVOLVER wired for firmware and pumps connected. The third pump is not included in this image, but is used during the recording of demonstration videos.

**Figure 13.**
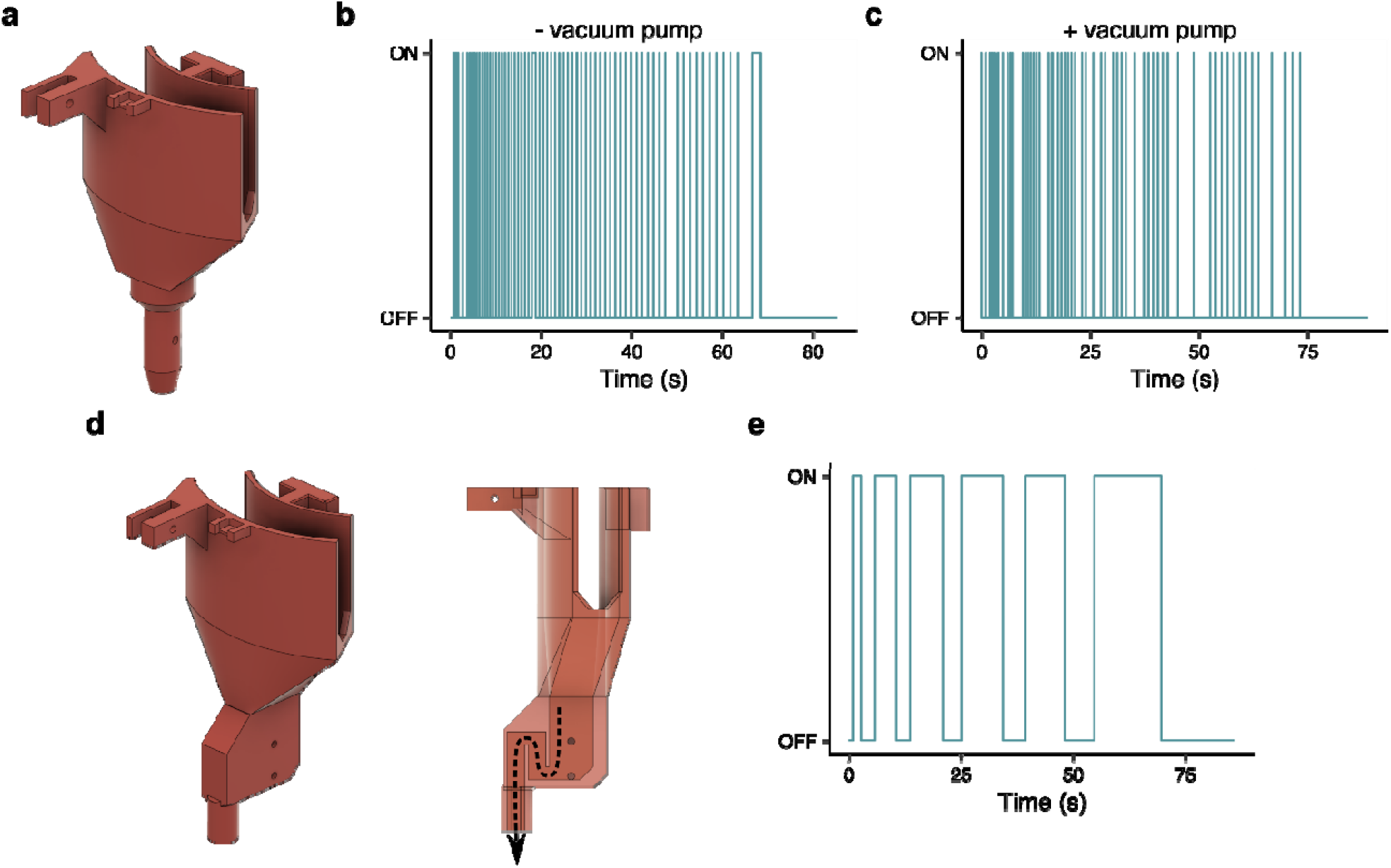
Design and characterization of the waste sensor. (a) Waste collector version 1, (b) Detection of liqui drops without a third pump attached to the waste collection version 1 and (c) with a third pump attached, (d) Waste collector version 2, (e) detection of self-flushing mechanism where no vacuum is required for waste collector version 2. Note: waste collector version is only discussed in Section 7.1. All other sections refer to waste collector version 1.

**Figure 14.**
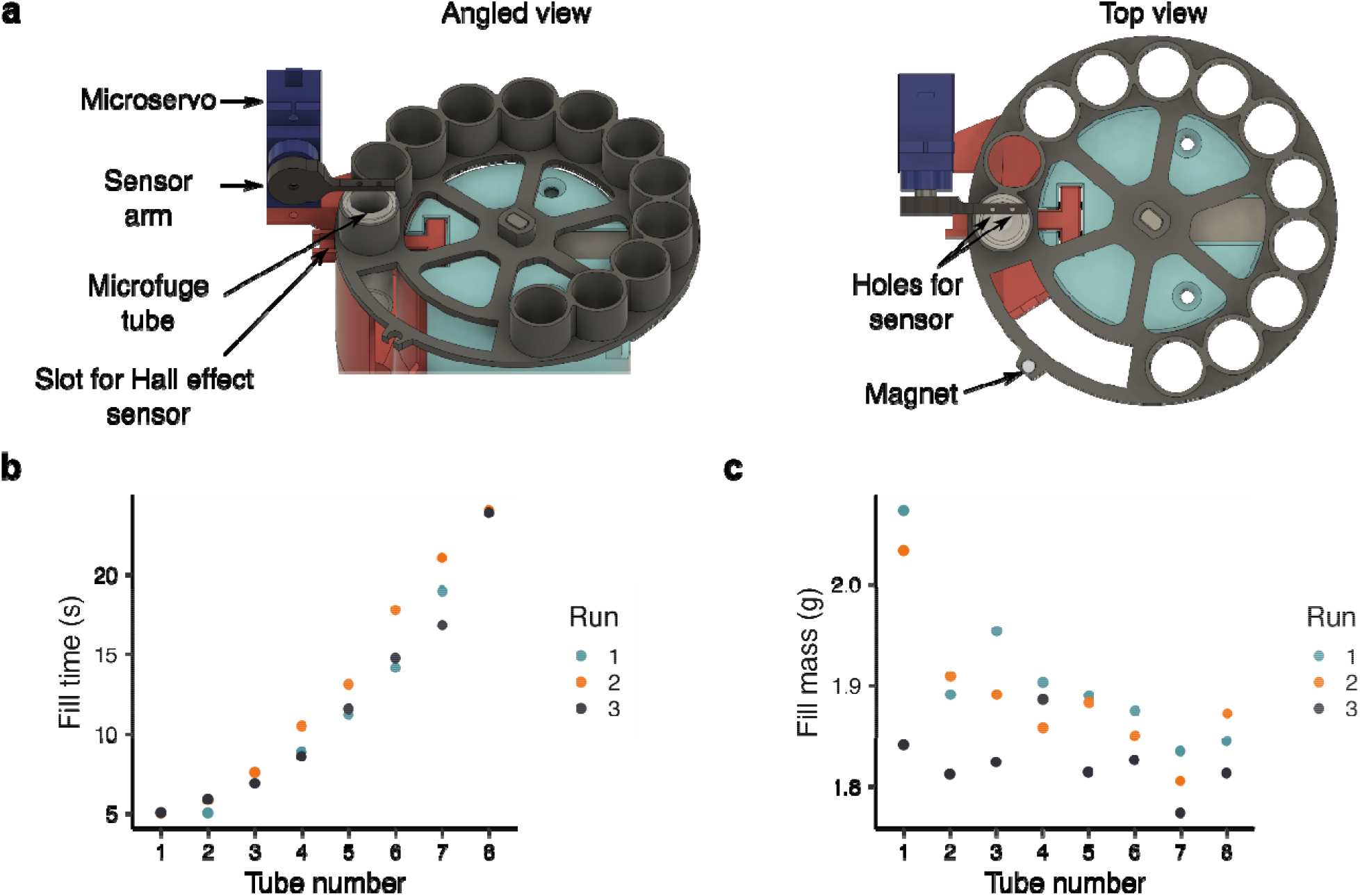
Design and characterization of the tube sensor. (a) Detailed view of the tube sensor component, (b) Time to fill and (c) weight of each tube after adding 16 mL of buffer to the column.

#### 2.1.2. Closed-loop functionality

Second, REVOLVER boasts closed-loop features to address several inconveniences that come with manual protein purification using gravity columns. Extensively used columns can have slower flow rates than others due to poor column filter cleaning, or the presence of solid precipitates inside of the filter that block the passage of the solution. Even clean columns can have minor variations in flow rates based on how well the resin is packed and how much liquid is in the column (*i*.*e*., hydrostatic pressure). This means that the time spent in each operation is not always known beforehand. Consequently, the user must be vigilant in watching the column to ensure the resin does not dry during the wash steps, or that each fraction of many has finished eluting without overflowing and spilling precious sample. This can be an inefficient use of time and limits the number of samples to be processed at once depending on the skill and focus of the operator.

One alternative to address this issue was to have a peristaltic pump system to ensure a constant flow rate through the resin like in FPLC systems, which would allow for open-loop operation of the system since the time required for each operation can be predetermined. However, this would complicate the design and require expensive equipment. Another alternative was to attach a flow rate sensor to the exit hole of the column to calculate the amount of solution that had passed through the resin, yet these sensors are not cheap (>$50 CAD), inaccessible for low flow rates (< 1 mL/min), and require custom fittings to adapt to columns. Rather than attempting to measure low flow rates, we designed a simple sensor to determine whether a conductive solution was flowing through the system. Specifically, we used a pair of pins that were physically close to one another (~1-2 mm) but not touching, one connected to ground and the other to a digital input pin in the Arduino. The pair of pins act as a digital-like switch when there is liquid present between them to carry the current across, which is detectable by the firmware.

Two pairs of these liquid-detecting pins exist in REVOLVER: one pair inside the waste collector to monitor solution (*i*.*e*., cell lysate flow-through and wash) that would pass through the resin into waste, and the second mounted onto the servo motor to detect when each of the microfuge tubes were filled during the final elution step. The servo lifts the pins out of the microfuge tube to allow the plate to rotate and position the subsequent tube, and then lowers the pins to monitor the level of liquid again. Because all solutions regularly handled during protein purification were readily conductive (300 – 600 mM NaCl), our liquid sensors (the pair of pins) were effective at detecting the movement of solutions across the resin. Sensor testing is detailed in Section 7.1 and 7.2 (Fig. 15-17).

**Figure 15.**
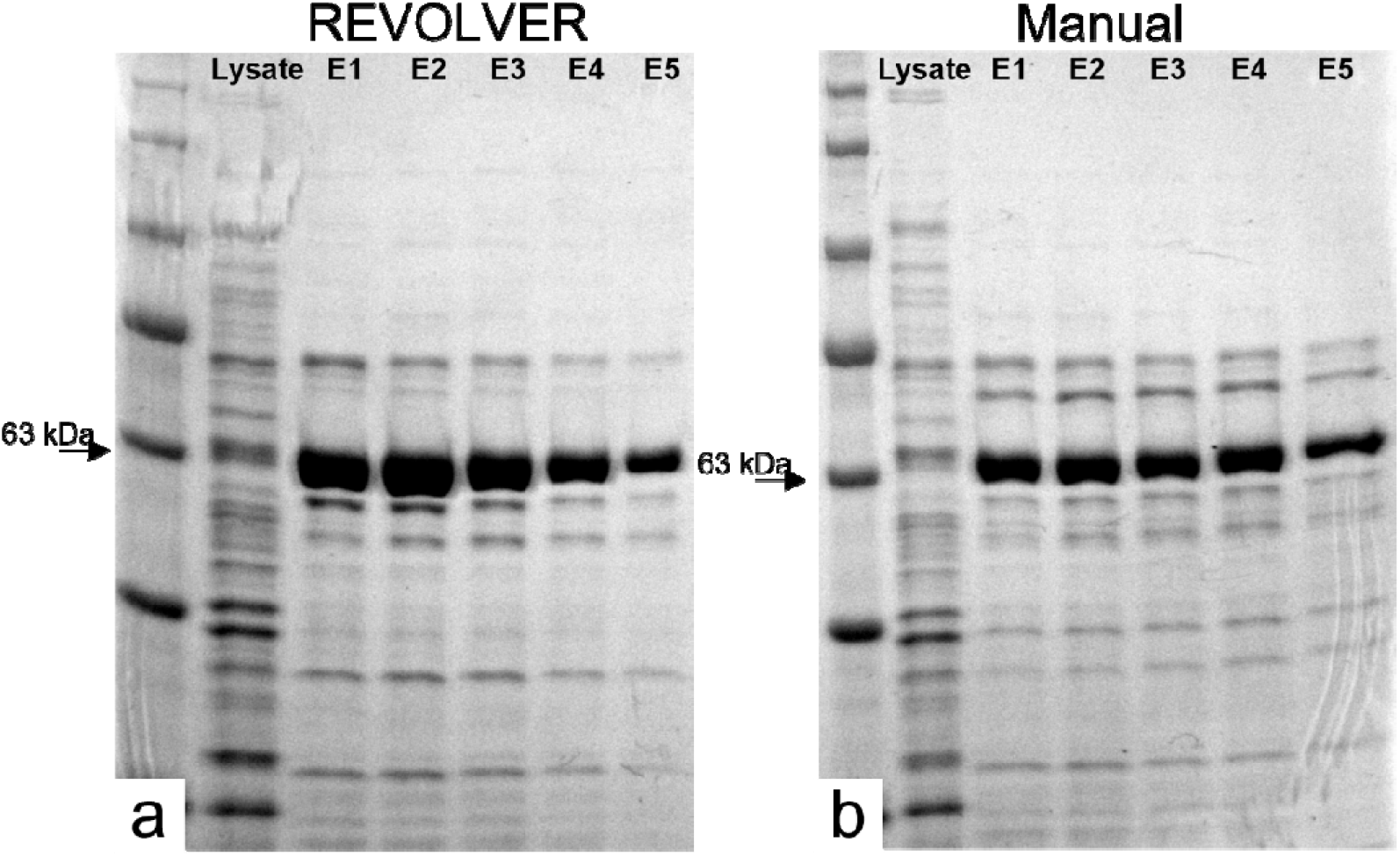
SDS-PAGE analysis of *Cc*SBPII protein purification. using (a) an automated protocol using REVOLVER, and (b) manual protocol by hand.

#### 2.1.3. Flow reduction to minimize resin bed disturbance

Third, REVOLVER includes a hardware feature to reduce the flow of solutions into the column. High flow rates (> 0.5 L/min) directly entering the column as a single stream can lift the packed resin from the base and expose the column’s filter. This results in the solution following the path of least resistance and avoiding paths through the resin; thus reducing protein binding, wash stringency, and target protein yields. We avoid high flow rates by first supplying the DC motor pumps with the minimum voltage required (6V, a setting on the power source). We then use a 3D-printed adaptor we designed for the column’s entrance hole that dissipates the force of the solution entering the column and evenly distributes it onto the interior column wall to gently fill the column. This piece is called the flow diffuser (Fig. 2c).

### 2.2. MULTI-VOLVER: multiple sample fraction collector

The objective of the MULTI-VOLVER (Fig. 3a) is to take multiple cell lysates as inputs, and return multiple sets of purified proteins in microfuge tubes as outputs with no human intervention. One could manufacture multiple REVOLVERs each with their own pumps and power supplies for automated parallel protein purification. These components represent the largest fraction of the cost of REVOLVER, yet the pumps remain idle during most of the protein purification and the capacity of the power supply is underutilized. The MULTI-VOLVER is a cheaper solution that we devised to economize the use of a single set of pumps and one power socket instead. To be exact, six REVOLVERs with their own pumps and power supplies would cost $990 CAD. The MULTI-VOLVER brings this price down to $310 CAD – a 60% reduction for the same throughput.

This is accomplished through an additional central distributor device controlled by an Arduino that can service the “needs” of up to six REVOLVERs (*i*.*e*. the Workers) (Fig. 3b and 3c). “needs” here referring to completing a wash step and being ready for another round of wash to be pumped into the column, or for the elution of the protein to begin. The Distributor services the Workers through two additional functionalities: 1) an (optional) auto-homing mechanism that allows a distribution arm to find the location of the gravity columns, and 2) a distribution arm that rotates in circular fashion to guide the ends of the pumps towards the gravity column that is ready for the next solution to be pumped inside. Moreover, the Distributor coordinates the individual REVOLVERs and passes the instructions for executing the purification protocol; this communication protocol is detailed in Section 2.4. Since the MULTI-VOLVER is designed to have up to six REVOLVERs working together, arranged symmetrically (Fig. 3b), it benefits from the three design features previously described (Section 2.1.1. to 2.1.3.). Assembly details are described in Section 5.0.

#### 2.2.1. Auto-homing to locate gravity columns

Unlike the REVOLVER (Fig. 2a), the lip of the gravity columns in the MULTI-VOLVER can be fitted with a special ring holding a Hall sensor that can be attached with hot glue. This allows the MULTI-VOLVER to automatically detect the position of the Workers’ columns (Fig. 3a). This ring can then be slid up and down such that it is beneath the distribution arm’s magnet when the distribution arm is rotated. When the position of the magnet is aligned directly on top of the Hall sensor during the auto-homing routine, the Distributor and Worker specific for that Hall sensor communicate to record the angular position of the distribution arm. This essentially provides the Distributor with the angular “address” for that Worker so in the future when solution is ready to be pumped into that Worker’s column, the Distributor knows which address to move the distributor arm to. It is also possible to manually rotate the distributor arm to store the address for each Worker. Details for automatic and manual homing of the distributor arm is further described in Section 6.1.

#### 2.2.2. Distribution arm to dispense solutions

Since there is only one set of pumps, which includes one for the wash solution and the other for the elution solution, the Distributor has to move the distributor arm to the Workers’ addresses for the pumps to deliver the wash/elution solution to the right column. The distributor arm is controlled by a stepper motor, which is controlled by the Arduino that is housed inside the electronics box of the Distributor (Fig. 3c). To prevent tangling of the tubes for the pumps, the firmware restricts the distributor arm from making a full 360° rotation. More specifically, the firmware determines a set point where the distributor arm cannot cross (*e*.*g*., if the distributor arm is at Worker 1, and the Distributor needs to pump solution into left-adjacent Worker 6, and the set point is between Worker 1 and 6, it will rotate counter-clockwise ~300° rather than clockwise 60°). The set point is automatically determined based on the distributor arm’s position when the MULTI-VOLVER is turned on.

### 2.3. Simplified Wiring – PCB Design

REVOLVER uses an Arduino Nano as a microcontroller but there are no off-the-shelf expansion boards to connect it to the components in our system. We therefore designed a custom printed circuit board (PCB) that would fit in the electronic boxes of REVOLVER. We ordered the PCB through JLCPB, although any other PCB manufacturer should be able to produce it.

The headers, sockets, and other electrical components (*e*.*g*., resistors, capacitors, diodes, transistors, screw block) were soldered on according to the system configuration (Fig. 6, Supp.Fig. 3). When completed, the PCB allows the user to simply plug the different components (*e*.*g*., motors and sensors) into the PCB for easy assembly of the hardware and electronics. The PCB includes extra headers for a 5V line that can be controlled by the remaining analog and digital pins we did not use. Another line set to be the same voltage delivered by the power supply is set to is also provided (*e*.*g*., 1 – 20V using our power supply). These were included in case the user wishes to modify the systems and power other useful features. While the PCB is a convenient feature of our design, it is not fully required and complete wiring diagrams can be found in Supp.Fig. 4.

### 2.4. Firmware and Communication Protocols

A requirement of our system was that it should be flexible enough to handle different purification protocols without having to update the firmware in each microcontroller. To achieve this, we made REVOLVER programmable by designing firmware that can take instructions and parameters from the user, and then execute them based on a predefined set of operations (*e*.*g*., wash, fill tubes).

The user interface with the REVOLVER is via serial USB connection with the user’s personal computer, specifically through the <u>Arduino IDE</u> software (Fig. 4a). Similarly, the user interface with the MULTI-VOLVER is via serial USB connection with the Arduino IDE on PC, except the connection exists with only the Distributor, not the Workers. The Workers are connected to the Distributor via an inter-integrated circuit (I2C) bus instead (Fig. 4b). In all cases each Arduino Nano microcontroller has to be uploaded (flashed) with the corresponding program for the REVOLVER and the MULTI-VOLVER’s Distributor and Workers before this serial USB/ serial USB+I2C bus user interface can be used, respectively.

For the Arduino Nano microcontroller configured for an independent REVOLVER, Fig. 4c depicts the main algorithm that takes inputs from the user through the Arduino IDE’s serial monitor. User commands take the form of “<?>” where the “<” and “>” tell the program that a command is being made, and the “?” between the < > symbols represents a capitalized letter corresponding to a specific command the user wishes to run, plus the required arguments for that command. The firmware then executes the instruction based on a preset list of operations. A sequence of commands can be made using the notation in sequential form <H><P><C>, for example. Table 1 contains a list of commands that can be currently interpreted and executed by the hardware.

**Table 1.** List of commands for the REVOLVER and MULTI-VOLVER system.

| Task | Input | Argument(s) | Example | Interpretation |
| --- | --- | --- | --- | --- |
| **Tasks below can be used for both the REVOLVER and MULTI-VOLVER** |  |  |  |  |
| Auto-home | H | n/a | <H> | Home REVOLVER |
| Rotate plate<br>(REVOLVER Only) | R | Number of steps,<br>Direction (0 or 1) | <R,1000,1> | Rotate motor 1000 steps |
| Pump solution | P | Pump volume (mL),<br>Pump ID | <P,20,1> | Turn pump on for 20 mL<br>from pump ID 1 |
| Collect waste | C | n/a | <C> | Wait and collect waste |
| Fill tubes | F | Number of tubes | <F,3> | Fill three tubes |
| **Tasks below are specific for the MULTI-VOLVER** |  |  |  |  |
| Locate Workers | L | n/a | <L> | Determine columns'<br>angular positions |
| Rotate plate<br>(MULTI-VOLVER only) | R | Number of steps,<br>Direction (0 or 1), I2C<br>device address | <R,1000,1,1> | Rotate motor of Worker<br>device #1 by 1000 steps* |
| Manual address finding | S | I2C address | <S,1> | Store the angular<br>position of Worker #1 |
| Rotate distributor to visit<br>a Worker | V | Device | <V,1> | Go to Worker #1 |
| Send instructions to<br>Workers | X | *input functions*<br>*between* | <X><H><P,10,1><C><X> | Home, Pump 10 mL from<br>pump ID 1, collect |

Building on this algorithm in Fig. 4d, the Arduino Nano microcontroller configured for the MULTI-VOLVER’s, the Distributor device also reads and parses instructions sent via USB. To run a full protocol, the Distributor first identifies the locations of the Workers (*cf*. Section 2.2.1.). The user then inputs a command sequence through serial monitor to the Distributor, which is then sent from the Distributor to the Workers one command in the sequence at a time via the I2C bus. To do so, the user must first enclose the desired commands between <X> commands; this notifies the Distributor to store the instructions. The Distributor continuously checks for whether a Worker has completed its singular command, which if True (*i*.*e*., completed), the next command in the sequence is sent until there are no more commands for the Worker. For the Arduino Nano microcontrollers configured as Workers, they sit idle until commands are sent to them from the Distributor. However, they idle for a very short period of time because the Distributor is continuously checking for whether a Worker’s task is done. Once all Workers finish all tasks stored by the Distributor, the whole system stands idle and can accept a new set of instructions without being reset. Different command sequences given to different workers for the MULTI-VOLVER described in this paper is currently not supported. To summarize by analogy, the Distributor has the plan and knows when to tell the Workers to perform their next step until they are done. The Workers simply wait to be told what to do after completing a command.

While we designed the MULTI-VOLVER to have up to six REVOLVERs networked together, I2C bus with Arduino Nano microcontrollers can theoretically allow a modified MULTI-VOLVER to control 127 Workers, which is five-times more than The Protein Maker [10]. Note, however, that building such a large MULTI-VOLVER with the current configuration of a central distributor would not be practical. However, the design can be adapted by incorporating a decentralized distributor that can be used to scale up to use the control capacity of the Arduino microcontrollers.

### 2.5. Fraction collector comparison

While other open-source fraction collectors have been published, such as the colosseum [12], our design goes beyond fraction collection and provides a complete platform for automatic protein purification from start to end. Our design differs from the colosseum in two ways. First, we prioritized shorter print times and less material usage. The colosseum device requires over 500 g of filament, with a total print time of >50 hrs. Our REVOLVER requires less than 150 g of filament with a total print time of less than 8 hours with the recommended settings. The MULTI-VOLVER requires roughly the same amount of filament and print time as the colosseum, yet it separately collects fractions for up to six different protein samples.

Second, the colosseum is an open-loop system that requires precise calibration of the pumps and flow rates to know how much liquid to dispense in each fraction. Our systems are closed-loop and are thus amenable to non-pressurized gravity column-based workflows, which simplifies the work the user needs to do. Nonetheless, our devices are limited in how many fractions they can collect (12 fraction/REVOLVER). They also require custom PCBs rather than commercial CNC shields like in the colosseum, meaning that the user does need to order PCBs, and then solder the required components onto it. Nonetheless, the systems we present here are more than just fraction collectors. To summarize our device:

- We designed a **low-cost** and **scalable** system for automatic parallel protein purification through gravity column-based workflows.
- The REVOLVER and MULTI-VOLVER are **closed-loop** systems that allow the user to load a cell lysate, leave the system unattended, and return to purified proteins in small fraction for SDS-PAGE analysis.
- We provide all files necessary for modifying everything about these devices to meet the users’ needs. This includes the hardware and firmware that is **modular** and **customizable**.
- Our systems have a **small form factor** the allow them to be stored in cold rooms and refrigerators for proteins requiring colder handling protocols.

## 3. Bill of materials

A complete Bill of Materials can be found in Supp.Table 1.

## 4. Design files

REVOLVER

- a1, a2, a3 are combined into the upper-half of the REVOLVER (a2 is attached to the stepper that is attached to a1)
- a4 can replace a3 if the user wishes to use the siphon model of the waste collector (See Section 7.1. for details)
- a5 is attached to the servo that is attached to a3 (or a4)
- a6 and a7 are combined to form the electronics box, which is the lower-half of the REVOLVER that can be combined with the upper-half (a1, a2, a3)
- a8 is placed into the gravity column
- a9 (optional) is attached to the latches of a7 to create a gripper for the gravity column

MULTI-VOLVER

- b1, b2, and b3 are attached to the top and bottom of the PVC pipe (b3 and b11 should be slid onto the pipe before attaching b1 or b2)
- b4 and b5 are mounted onto the pipe using b3
- b6 is attached to the stepper that is attached to b2
- b7 is fitted onto the outer end of b6
- b8 is used to attach the REVOLVER’s latches (a7) to the latches on b1
- b9 is used to attach the REVOLVER’s latches (a7) to each others’ latches (a7)
- b10 is clipped onto the gravity column
- b11 is slid onto the PVC (see above for b1, b2, and b3) to replace a9
- b12 can be used to clips two b11’s together

ELECTRONICS

- c1 is used to connect all electronics components to the Arduino Nano microcontroller

## 5. Build instructions

This set of build instructions is divided into the REVOLVER system (Section 5.1), and the MULTI-VOLVER system (Section 5.2). Unique parts are shown in Figure 5a. Three composite parts are described in Supp.Fig. 1-3: the DC pump with tube adaptors, the Hall effect sensor board, and the soldered PCB. In particular, Supp.Fig 3 details how the headers, sockets, and electrical components should be arranged. In these instructions, we refer to the parts according to their numbering in Fig. 5 and their designators as described in the Bill of Materials (Supp.Table 1). For the 3D-printed MULTI-VOLVER parts, their names according to Table 2 are used.

**Table 2.** Design Files.

| ID | Name | Type | Open-source license | Location |
| --- | --- | --- | --- | --- |
| **REVOLVER** |  |  |  |  |
| a1 | revolver_tower | f3d, stl | BSD-2 | Available in repository |
| a2 | revolver_plate | f3d, stl | BSD-2 | Available in repository |
| a3 | revolver_waste_collector | f3d, stl | BSD-2 | Available in repository |
| a4 | revolver_waste_collector_syphon | f3d, stl | BSD-2 | Available in repository |
| a5 | revolver_servo_arm | f3d, stl | BSD-2 | Available in repository |
| a6 | revolver_box_top | f3d, stl | BSD-2 | Available in repository |
| a7 | revolver_box_bottom | f3d, stl | BSD-2 | Available in repository |
| a8 | revolver_flow_distributor | f3d, stl | BSD-2 | Available in repository |
| a9 | revolver_column_grip | f3d, stl | BSD-2 | Available in repository |
| **MULTI-VOLVER** |  |  |  |  |
| b1 | multivolver_base | f3d, stl | BSD-2 | Available in repository |
| b2 | multivolver_top | f3d, stl | BSD-2 | Available in repository |
| b3 | multivolver_box_mount | f3d, stl | BSD-2 | Available in repository |
| b4 | multivolver_box_bottom | f3d, stl | BSD-2 | Available in repository |
| b5 | multivolver_box_top | f3d, stl | BSD-2 | Available in repository |
| b6 | multivolver_distributor_arm | f3d, stl | BSD-2 | Available in repository |
| b7 | multivolver_arm_head | f3d, stl | BSD-2 | Available in repository |
| b8 | multivolver_linker_short | f3d, stl | BSD-2 | Available in repository |
| b9 | multivolver_linker_long | f3d, stl | BSD-2 | Available in repository |
| b10 | multivolver_hall_ring | f3d, stl | BSD-2 | Available in repository |
| b11 | multivolver_column_gripper | f3d, stl | BSD-2 | Available in repository |
| b12 | multivolver_gripper_post | f3d, stl | BSD-2 | Available in repository |
| **ELECTRONICS** |  |  |  |  |
| c1 | revolver_PCB | grb | BSD-2 | Available in repository |

Suggested 3D-printing orientations, as well as filament weight and time estimates for the prints, are summarized in Supp.Fig. 6 for the REVOLVER and Supp.Fig. 7 for the MULTI-VOLVER.

### 5.1. REVOLVER Assembly

#### 5.1.1. Waste collector assembly

#### 5.1.2 Tube sensor assembly

#### 5.1.3. Motor tower assembly

#### 5.1.4. Full REVOLVER hardware assembly

#### 5.1.5. Wiring hardware with firmware

Step (a): pull the wiring of the servo (Part 20) through the top of the electronics box (Part 30)

Step (b): connect the stepper’s wiring (Part 19) to the STEPPER pins of the soldered PCB (Part 23). Usually the blue wire is oriented to be on the far left

Step (c): connect the servo’s wiring (Part 20) to the SERVO pins of the soldered PCB. Usually the brown wire is oriented to be on the far left

Step (d): attach three jumper (FF) wires (Part 18) to the pins of the Hall sensor (Part 25) attached to the waste collector (Part 29)

Step (e): connect the ends of the jumper (FF) wires from the previous step to the H1 pins of the soldered PCB. Take note that the H1 pins are not aligned in the same sequence as the Hall sensor pins

Step (f): attach two jumper (FF) wires to the TUBE pins

Step (g): attach two jumper (FF) wires to the WASTE pins

Step (h): attach two alligators (Part 14) to the PUMP1 pins, and another two alligators to the PUMP2 pins

#### 5.1.6. Connecting pumps to solutions

#### 5.1.7. Vacuum-drained waste collector (recommended, but optional)

We have found that the waste collector more reliably drains fluid when a third pump is constantly drawing solution out of it. A future iteration of our designs can modify the PCB to accommodate the control of a third a pump for this purpose.

Step (a): connect the negative pressure end of the pump (Part 8) to the small tubing piece from Section 5.1.6. step (f)

Step (b): connect the positive pressure end of the pump to another small tubing piece cut to the user’s desired length (recommended 45-60 cm) and place the other end of this tubing piece into a waste receptacle

Step (c): connect the pump to another power supply (Part 21) using a pair of alligators (Part 14)

### 5.2. MULTI-VOLVER Assembly

#### 5.2.1. Hardware assembly

The following steps assume the user has already created the number of individual REVOLVERs they wish to use with the MULTI-VOLVER. Refer to Section 5.1 for REVOLVER assembly, and Supp.Fig. 3 for the correct PCB configurations for the Distributor and Worker.

Step (a): slide two column grippers (b11) onto the 30 cm long PVC pipe, then slide the box mount (b3) below the column grippers. Attach the electronics box bottom (b4) to the box mount using 6 mm M3 screws (Part 4)

Step (b): align the column grippers such that the two slots for the small tubing pieces are directly on top of one another, then connect the column grippers together using six gripper posts (b12)

Step (c): attach the base (b1) below the box mount onto the end of the PVC pipe. Holes are provided if the user wishes to screw the components together, or hot-glue could be used instead

Step (d): use two 12 mm M3 screws (Part 3) and two M3 nuts (Part 5) to attach a stepper (Part 19) to the top (b2), similar to Section 5.1.3.

Step (e): attach the above component onto the other end of the PVC pipe, above the column grippers

Step (f): apply a small amount of hot-glue to the stepper’s axle, and connect the distributor arm (b6)

Step (g): insert a magnet (Part 24) into the hole of the distributor arm

Step (h): gravity columns can then be clipped in place using the column grippers

Step (i): similar to Section 5.1.1. steps (h-k), attach a Hall sensor (Part 25) to the slot in the Hall ring (b10)

Step (j): clip the Hall ring with the Hall sensor approximately 1 cm below the lip of the gravity column

Step (k): using short linkers (b8), connect the desired number of REVOLVERs (with their Arduino Nano microcontrollers configured as Workers) to the base (b1) as shown in Fig. 2. Workers that directly adjacent to each other can be connected using long linkers (b9)

Step (l): rotate the columns (Part 1) and the column gripper

#### 5.2.2. Wiring hardware with firmware

Step (a): attach the soldered PCB to the electronics box (b4) using 6 mm M3 screws (Part 4), then connect an Arduino Nano microcontroller to the soldered PCB for the Distributor configuration

Step (b): connect each column’s Hall sensor from Section 5.2.1. step (j) to jumper (FF) wires

Step (c): connect these jumper (FF) wires from the previous step to the H2 pins of the corresponding Worker’s PCB. Jumper (MM) wires may need to be used to cover the distance between the Hall sensor and the PCB

Step (d): link each Worker to the Distributor by connecting one SDA pin, one SCL pin, one “12V” pin and one GND pin of the Worker to the same set of pins on the Distributor using a combination of jumper (MM) and (FF) wires to cover the distance. For example in a 6-Worker MULTI-VOLVER, the Distributor will have six sets of SDA, SCL, “12V”, and GND pins occupied, each set connected to one of the six Workers. “12V” is in quotations because the actual voltage is set by the power supply

Step (e): attach two alligators (Part 14) to the PUMP1 pins, and another two alligators to the PUMP2 pins

Step (f): attach each pair of alligators to a pump’s positive and negative clips (Part 8)

Step (g): loosen the screw block of the PCB and insert one jumper (MM) wire (Part 17) into each positive and negative terminal, then re-tighten the screw block to lock the wires in place

Step (h): connect the ends of the jumper (MM) wires from the previous step to the power supply’s adaptor piece, then connect the adaptor to the power supply (Part 21)

#### 5.2.3. Connecting pumps to solutions

Step (a): perform Section 5.1.6. steps (a-f) with both pumps (Part 8) connected to the Distributor (not the Workers) and using small tubing pieces cut to approximately 80-100 cm

Step (b): clip the 1-way valve (Part 11) for each pump into the arm head (b7)

Step (c): apply hot glue into the slot on the opposite side of the 1-way valve slots, and insert the distal end of the distributor arm (b6) from Section 5.2.1. step (f) into the slot

Step (d): insert the small tubing pieces from step (a) into the slots aligned earlier in Section 5.2.1. step (b)

Step (e): repeat Section 5.1.7. steps (a-c) use Y connectors (not shown in Fig. 5, but included in Supp.Table 1) and small tubing pieces (cut to 30-50 cm) to connect the Workers drainage tubing (Fig. 11f) to the negative pressure end of a third pump to help drain all Workers.

## 6. Operation instructions

### 6.1. Set-up

Prior to initializing the REVOLVER or MULTI-VOLVER systems, the user must upload the correct program into the Arduino Nano microcontroller (termed “flashing”). These programs can be found in the following links to our GitHub repository: <u>REVOLVER</u>, <u>Distributor</u> (MULTI-VOLVER), and <u>Worker</u> (MULTI-VOLVER). The Workers’ microcontrollers must each be assigned a unique integer under the code *const byte I2Caddress* before flashing. The values must be in the range of 1 – 128 (inclusive); we recommend labelling them 1 to 6 for a full 6-device MULTI-VOLVER (though it is not mandatory to label in this sequence as long as the user knows which Worker has been assigned which address). In contrast, the programs for the REVOLVER and Distributor can be flashed without immediately changing any variables, but will very likely require the user to modify a few variables after the initial flash, as detailed in Section 6.2 due to differences in DC pump manufacturing. Once all variables are set for each configuration and flashed onto the appropriate system, the user does not need to re-flash the microcontroller every time they use it as long as the addresses and pumps stay the same.

#### 6.1.1. Initializing REVOLVER

The REVOLVER is initialized by first connecting the Arduino Nano microcontroller configured for the single device to the user’s PC via serial USB. Once the program is flashed onto the microcontroller through the Arduino IDE, the user should open the serial monitor to be able to communicate with the REVOLVER. To test the connectivity of the various components and check whether they were assembled correctly before using the REVOLVER to purify a protein, we have created a checklist of tasks that allow the user to identify and address problems beforehand (Table 3). While we have tested the build extensively with the parts obtained with our printers, the user should expect to encounter *some* of the common challenges; the solutions we have offered should be considered different techniques to optimize the REVOLVER system’s performance.

**Table 3.**
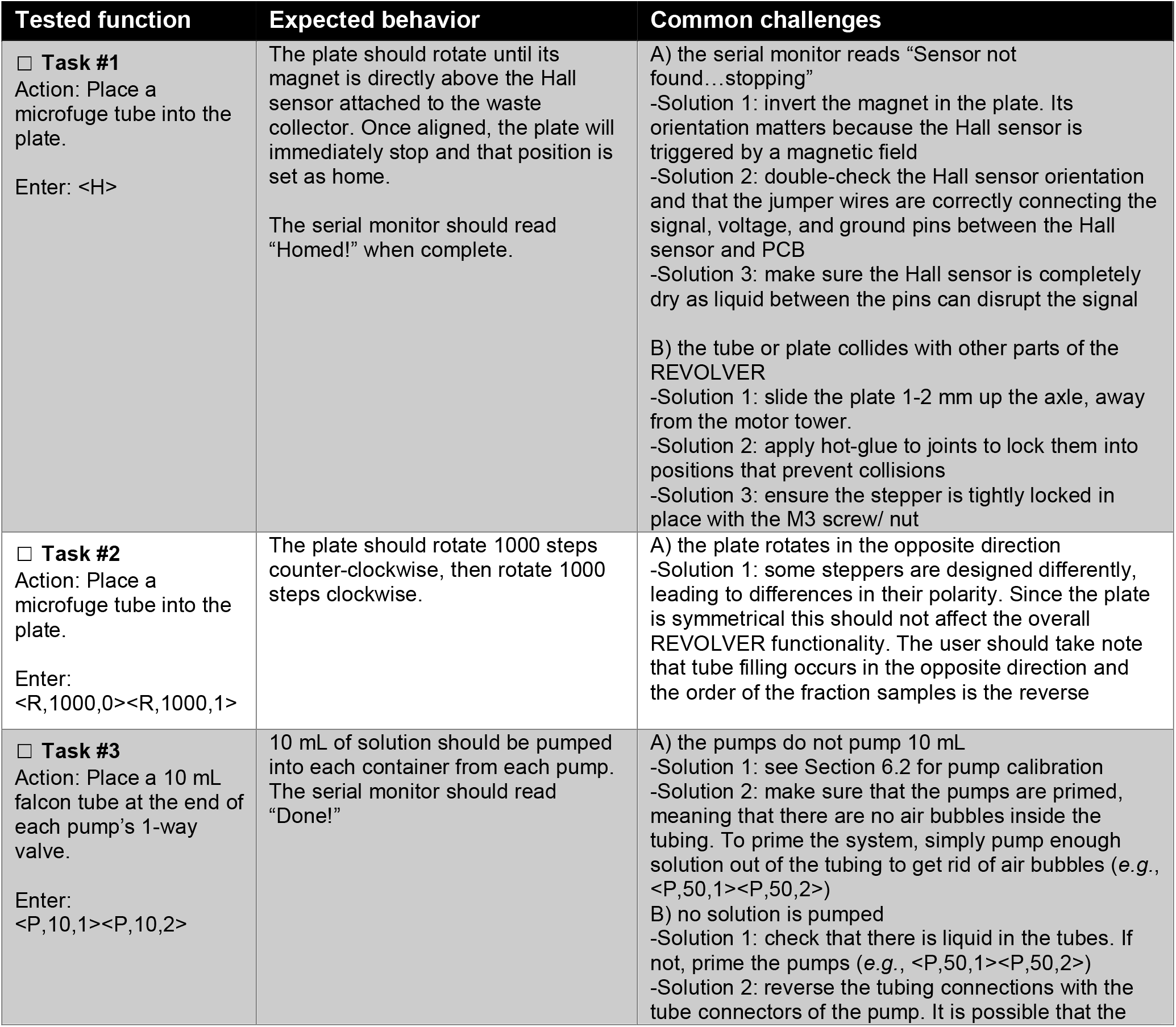
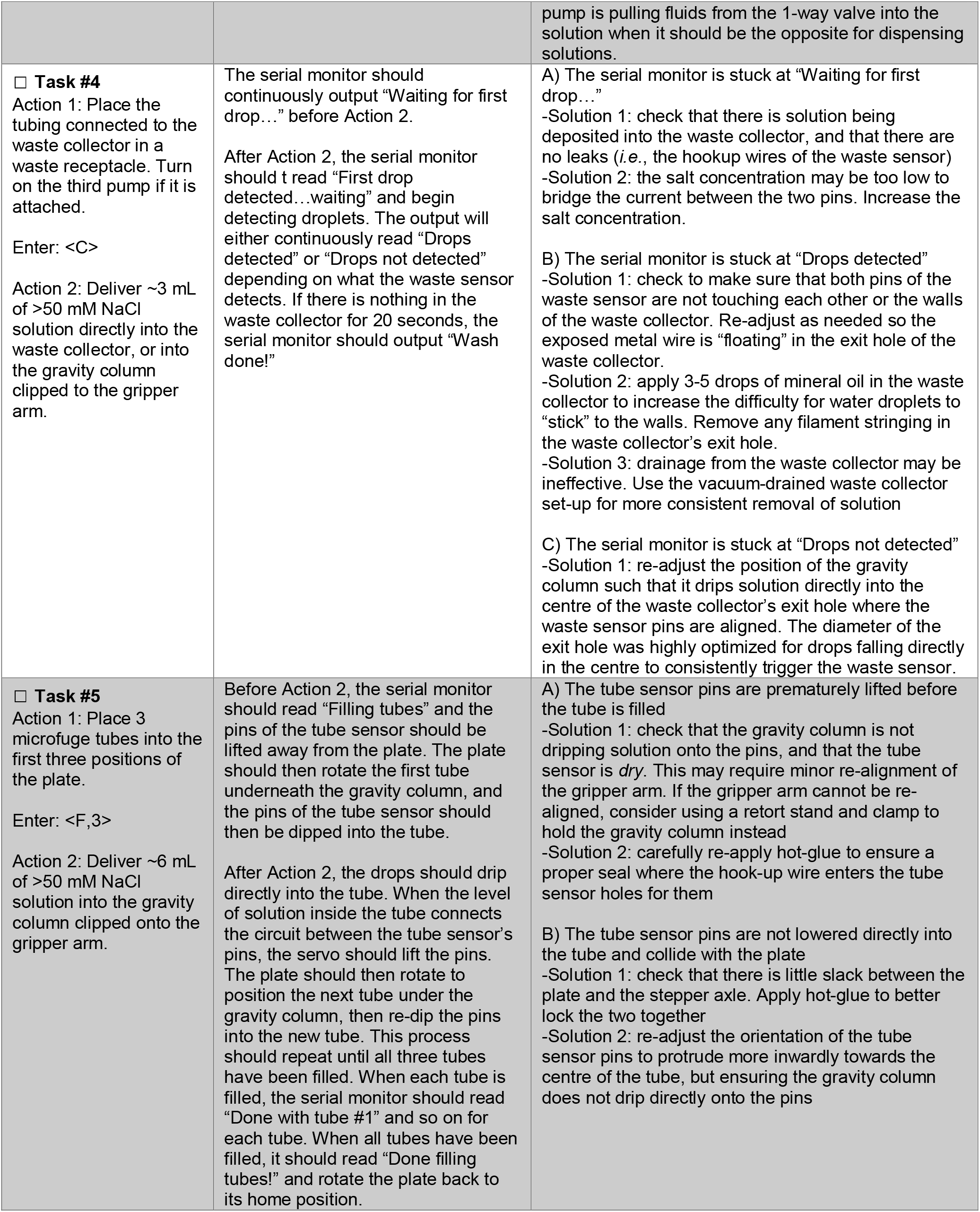
Checklist for initialization of the REVOLVER system.

**Table 4.** Checklist for initialization of the MULTI-VOLVER system.

| Tested function | Expected behavior | Common challenges |
| --- | --- | --- |
| ☐ **Task #1**<br>Connect Distributor to PC via serial USB and open the serial monitor on the Arduino IDE | The serial monitor should read "Scanning I2C bus..." at first, then read "I2C device found at address: 1" and repeat the corresponding message for all other Workers connected to the Distributor. When complete, it should read "Found X devices where X is the number of Workers. Lastly, it will prompt the user with the message "Setup done...please locate devices". | A) one of the Workers is not detected<br>-Solution 1: check to make sure the jumper wires connected the SDA/SCL/GND/"12V" are not wet and therefore connected to each other via liquid<br>-Solution 2: user did not change the unique I2C address of each Worker before uploading the program to the microcontroller. There is a special const byte for this at the top of the program |
| ☐ **Task #2**<br>Action: Place a microfuge tube into the plate for all Workers<br>Enter: <X><H><X> | The plates should rotate until their magnet is directly above the Hall sensors attached to the waste collectors. Once aligned, the plates will immediately stop and those positions will be set as home. | See Table 3, Task #1 – Common Challenges. |
| ☐ **Task #3**<br>Action 1: Place a microfuge tube into the plate.<br>Action 2: Attached pump tubing to distributor arm using the arm head and 1-way valves.<br>Enter:<br><R,1000,0,0><br><R,1000,1,0><br><R,1000,0,1><br><R,1000,1,1> | The distributor arm will rotate 1000 steps clockwise, then 1000 steps counter-clockwise. Next, the plate for the Worker at address 1 will rotate 1000 steps clockwise, then 1000 steps counter-clockwise. | A) the distributor arm or the plates rotate in the opposite direction as intended (0 = clockwise, 1 = counter-clockwise)<br>-Solution 1: some steppers are designed differently, leading to differences in their polarity. Since the plate is symmetrical this should not affect the Worker's functionality. The user should take note that tube filling occurs in the opposite direction and the order of the fraction samples is the reverse.<br>This may however affect the distributor arm's functionality. It is recommended that the user purchase steppers that all utilize the same direction for 0 and 1, and work with the system in the opposite direction |
| ☐ **Task #4**<br>Enter: <L> | The serial monitor should read "Finding locations of devices..." and the distributor arm should rotate counter-clockwise. Each time the magnet of the distributor arm aligns with the Hall sensor clipped onto each column and wired to a corresponding Worker, the serial monitor should read "Location of I2C #X found at n = Y" where X is the Worker's address and Y | A) one of the Workers is not detected<br>-Solution 1: check the orientation of the magnet and the Hall sensors; re-adjust as needed<br>-Solution 2: ensure the clipped Hall sensor and the distributor arm's magnet are as close as possible when the distributor arm is rotating<br>-Solution 3: user did not change the unique I2C address of each Worker before uploading the program to the microcontroller. There is a special const byte for this at the top of the program |

#### 6.1.2. Initializing MULTI-VOLVER

Similar to the REVOLVER, the MULTI-VOLVER is initialized by first connecting its Arduino Nano microcontrollers to the user’s PC via serial USB. Once the appropriate programs are loaded in the microcontrollers for the Distributor and Workers, the Distributor should be connected to the user’s PC followed by opening the serial monitor on the Arduino IDE to communicate with the MULTI-VOLVER system. Since the MULTI-VOLVER is comprised of a network of REVOLVERs, Table 3 is useful for optimizing the performance of the Worker. However, the MULTI-VOLVER has unique functions with their own tests for connectivity and proper assembly, as well as challenges. The common challenges should be expected, and the solutions we have offered should be considered different techniques to optimize the MULTI-VOLVER’s performance.

#### 6.1.3. Manually storing the Workers’ addresses for the MULTI-VOLVER

While the command <L> exists for auto-homing the distributor arm similar to how <H> auto-homes the plates, we’ve created an option that allows users to manually store the location (*viz*. angular position in steps) of the Workers’ columns. Below is the protocol:

Step (a): while the MULTI-VOLVER is off, rotate the distributor arm to a position directly opposite to where the pump tubing is clipped into the column grippers. When the MULTI-VOLVER is turned on, this step allows for the pump tubing in its neutral position to be step 0, which allows for the set-point of the distributor arm to best limit rotations that would tangle the pump tubing

Step (b): rotate the distributor arm using the <R> command until the 1-way valves of the arm head are sufficiently pointing into the column of the immediate left-adjacent worker (counter-clockwise)

Step (c): store this position using the <S> command and specify in the argument the Worker number Step (d): repeat step (b) and (c) above for the remaining Workers

Step (e): to test if the Distributor appropriately recorded the correct locations, use the <V> command with the specific Worker number in the argument to have the distributor arm move the 1-way valves to that Worker. The <P> command can be used to check if the solution is pumped into the column

### 6.2. Pump calibration

12V DC pumps have variable flowrates, so we suggest users calibrate their pumps by modifying four variables in the program for the Distributor’s Arduino Nano microcontroller and in the single REVOLVER firmware: smallVolumeCalibrationPump1, largeVolumeCalibrationPump1, smallVolumeCalibrationPump2, and largeVolumeCalibrationPump2. These are essentially calibration factors, which are found under the /* Constant properties of the hardware */ header. These variables distinguish between small volume (<= 1 mL) and large volume (> 1 mL) because the calibration required for either volume range is different (*viz*. the calibration is not linear at small volumes). The third (optional) pump does not need to be calibrated since it is only used as a constant vacuum to drain liquid from the waste collector. Below is the protocol:

Step (a): place both 1-way valves into a waste receptacle. Prime both pumps using distilled water before calibration by using the command <P,50,1><P,50,2> to remove air bubbles inside the tubing

Step (b): tare a scale for a 15 mL falcon tube, then place the tube underneath the 1-way valve for Pump 1 (the one connected to PUMP1 on the PCB)

Step (c): enter <P,1,1> to tell the pump to output 1 mL of solution from Pump 1. Step (d): measure the weight of the solution in the tube

Step (e): repeat step (b)

Step (f): enter <P,10,1> to tell the pump to output 10 mL of solution from Pump 1.

Step (g): repeat step (d)

Step (h): repeat steps (b-g), except this time for Pump 2. The user should have four measurements for the weight of the solutions at this point

Step (i): to calculate the new values for each variable, first calculate 1 divided by the weight of each solution pumped, then multiple this quotient by 1 mL or 10 mL for the small or large volume weight measurements, respectively. The products are the new value for the variables, respectively

Step (j): replace the default “1.00” for these variables with the new corresponding values

Step (k): repeat steps (b-h) to check if Pump 1 and Pump 2 accurately output volumes as entered

General guidelines for what calibration factors to use for a given output (measured volume) when 1 mL or 10 mL is inputted (desired volume) are described in Table 5. For example, if the user desires 10 mL from pump 1 and they measure 8.5 mL instead, they should assign the *const float largeVolumeCalibrationPump1* variable a value of 1.17. If the user desires 1 mL from pump 2 and they measure 1.3 mL instead, they should assign the *const float smallVolumeCalibrationPump2* variable a value of 0.76.

**Table 5.**
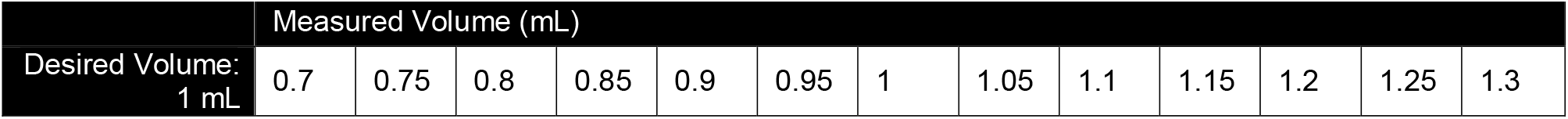
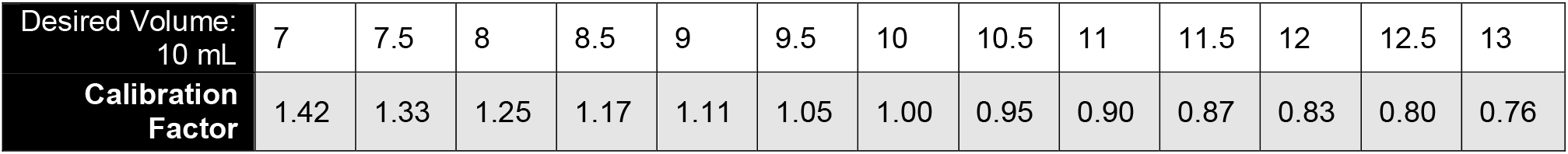
Illustrative Pump Calibration Factor Chart.

### 6.3. Running protocols

The REVOLVER and MULTI-VOLVER accept *command sequences*, which are lists of commands according to Table 1. To purify a protein, there are four primary commands: <H>, <P>, <C>, and <F>. Combinations of these commands mimic the same procedures a user would perform to wash affinity resins and elute target proteins with gravity columns. Table 6 describes typical command sequences for workflows that use a 30-mL gravity column for immobilized metal affinity chromatography. For the MULTI-VOLVER configuration, the Master can only execute commands <L>, <S>, <V>, and <R> on its own without having to send instructions to the Workers (*i*.*e*., without using the <X> command). We have also allowed the Master to pass the <R> command to the Workers outside of the main protocol in case the user wishes to perform manual homing of each REVOLVER.

**Table 6.**
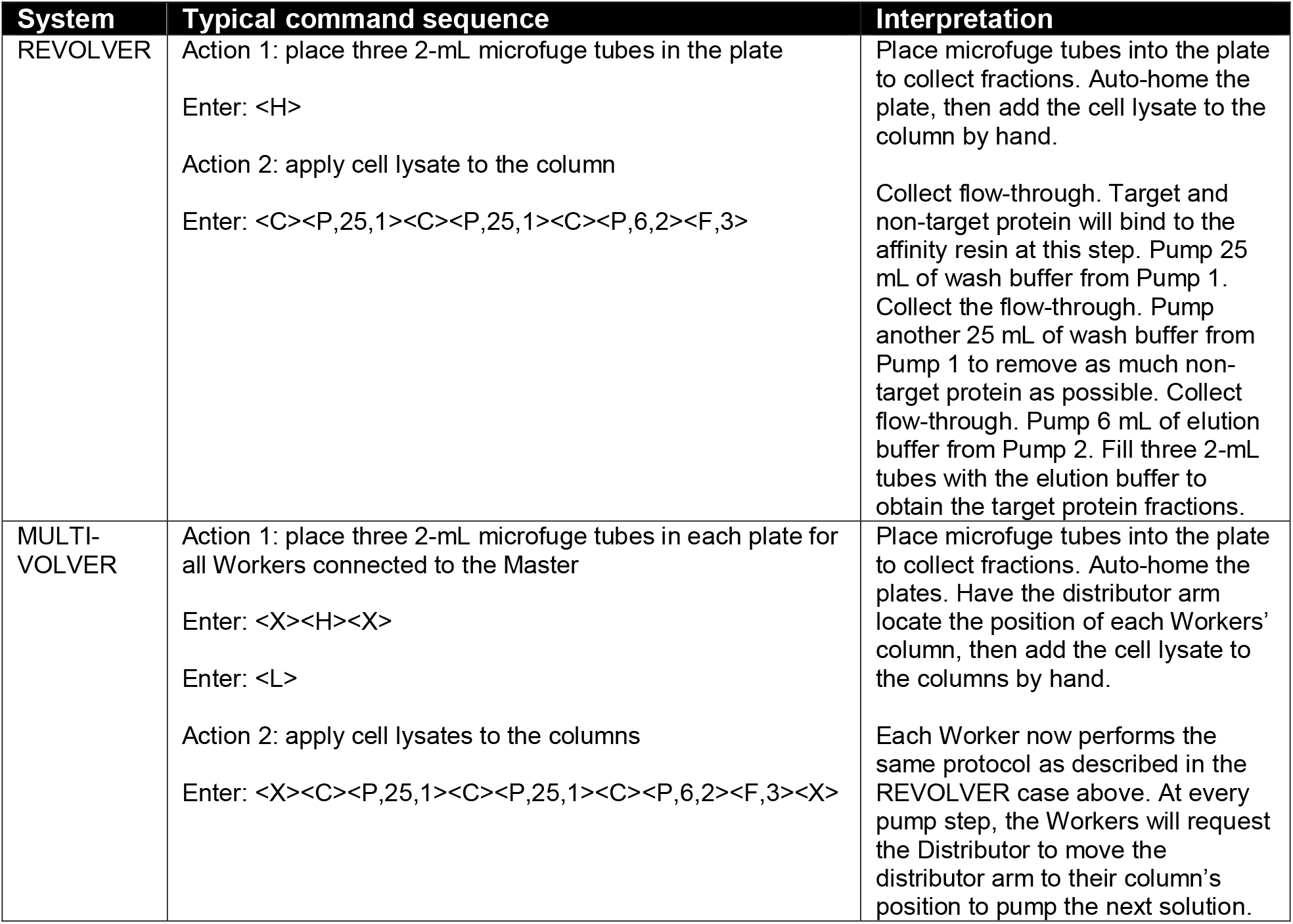
Example Protein purification protocols for the REVOLVER and MULTI-VOLVER systems.

The program expects the user to remember to add the cell lysate at the beginning of a protocol before the first <C> command. However, the program will notify the user to add lysate if no liquid is detected.

If the user is purifying a new protein for the first time and chooses to use either system for the task, it should be noted that there are parameters to optimize that cannot be controlled in the command sequences, which only control the wash volume, number of washes, and elution volume. These other parameters include the stringency of the wash solution and the strength of the elution buffer (*e*.*g*., concentration of imidazole for immobilized-metal affinity chromatography). For a detailed description of important purification parameters, we recommend the guide by the Structural Genomics Consortium *et al*. [2].

### 6.3. Clean-up and shutting down the systems

Since some of the REVOLVER and MULTI-VOLVER components make direct contact with biological samples (*i*.*e*., the waste collector) and due to splashing, they are prone to build-up of waste on surfaces. Below is a general protocol for cleaning the systems:

Step (a): remove protein fractions from the plate and resin from the column. Re-clip column back in place

Step (b): transfer tubing from wash and elution buffer solutions to a container of 20% ethanol

Step (c): enter <P,25,1> and allow the 20% ethanol to flow through. Repeat once more

Step (d): enter <P,25,2> and allow the 20% ethanol to flow through. Repeat once more

Step (e): unplug the power supplies, then the serial USB from the system

## 7. Validation and characterization

The REVOLVER and MULTI-VOLVER systems rely on the waste sensor and tube sensor functioning as expected (*viz*. they detect liquid when liquid is present, and detect nothing when liquid is absent). Section 7.1 and 7.2 test these sensors’ functionality, respectively. Section 7.3 describes our Supplementary Videos that show the REVOLVER and MULTI-VOLVER systems in action. Finally, Section 7.4 describes a real protein purification we performed manually and automatically with the REVOLVER.

### 7.1. Vacuum-drainage comparison

The waste collector (Fig. 13a) is comprised of a basin where the drops from the gravity column fall into, and an exit hole where the liquid is then drained. We created two small holes in this exit hole where pins (from the hookup wire) were inserted into. In the absence of the third pump to continuously drain from the exit hole, the two pins exhibit the detection behavior seen in Fig. 13b, where “ON” refers to liquid being detected by the sensor. Samples with a higher protein content can be more viscous and produce foaming in the collector, which reduces the draining efficiency and causes the sensor to stay “ON” even if the column has drained. Moreover, from our testing, the position of this tubing (*i*.*e*., laid flat vs. hanging off a bench) and its size (*i*.*e*., inner diameter) can affect drainage. Therefore, we recommend the user consider adding a third pump to the system. which improves the drainage of liquid from the waste collector leading to detection behavior seen in Fig. 13c.

As an alternative to using a third pump, we additionally designed a new waste collector (Fig. 13d) that had better drainage properties and does not require a third pump, thus reducing the cost of REVOLVER by roughly 20%. We were inspired by toilets (*viz*. siphons) and re-designed the exit hole to have an internal siphon that flushes itself. Rather than detecting the presence of droplets, the waste collector version 2 detects changes in the volume within the siphon. If there are no more droplets from the gravit column, the two pins will either stay connected or disconnected for a set threshold time, similar to the first version (Fig. 13a). Consequently, the detection behavior appears as Fig. 13e.

Since waste collector version 1 was initially designed and tested, this paper primarily focuses on it, however we encourage future users to consider first using waste collector version 2 since it does not require a third pump with its own power supply. Our testing shows version 1 with the third pump and version 2 both have consistent, reliable waste collector drainage. The programs for the REVOLVER and Distributor that we have provided (Section 6.1) can be used for both versions. The user simply needs to change the value assigned to the variable *const byte collectorVersion* from 1 (by default version 1) to 2 (version 2).

### 7.2. Tube filling consistency

As noted in Section 2.5, fraction collectors are often open-loop systems that rely on predictable flow-rates and pre-determined volumes to know how long to fill a collection tube before moving to the next fraction. However, gravity columns do not have predictable flow rates; the solutions flow through the column purely through gravity and pressure head. While one could pressurize the gravity column to control the flow rate of solutions, we deemed this approach too complex and expensive for an open-source solution. The tube sensor (Fig. 14a) follows the same principle as the waste sensor in that two metal pins detect the presence of liquid through the connection of a current. To demonstrate the importance of having our systems be closed-loop, Fig. 14b shows an experiment where we pumped 16 mL of elution buffer into the column and measured how long it took to fill each 2-mL microfuge tube in the plate. Since the gravity column’s flowrate changes, the tube sensor allows the REVOLVER and MULTI-VOLVER systems to relatively consistently output fractions of similar volumes without predetermined fill times per fraction (Fig. 14 c).

### 7.3. Protocol Demonstration

The details reported here are the result of several rounds of prototyping and testing. We recorded several videos to demonstrate the full functionality of the REVOLVER and MULTI-VOLVER. In Supp.Video 1, we demonstrate the REVOLVER performing a single wash step and elution into three microfuge tubes. In Supp.Video 2, we demonstrate the same protocol on three Workers networked to a Distributor in a MULTI-VOLVER system. The distributor arm’s auto-homing command <L> was shortly optimized thereafter and demonstrated in Supp.Video 3. While the I2C-based protocols we developed can easily handle the six Workers in a full MULTI-VOLVER, we showcased a 3-Worker version of the MULTI-VOLVER in Supp.Video 2. Nonetheless, we verified that it is possible to effectively drain liquid from six waste collectors with a single vacuum pump, as demonstrated in Supp.Video 4. The user should note that the addition of mineral oil to the waste collector prior to protein purification, which applies a hydrophobic coating onto the inner walls, aids in drainage to an appreciable degree (noted in Table 3. Task #4 Challenge B).

### 7.4. Purification of *Cc*SBPII, a nickel-binding protein

We wanted to confirm whether the REVOLVER system could actually purify a protein from a real-world cell lysate. A protein that is routinely purified in our work is *Cc*SBPII, a nickel-binding protein belonging to an ATP-binding cassette transporter from *Clostridium carboxidivorans*. To express the protein and process its cell lysate, the following protocol was followed:

Starter cultures for *Cc*SBPII were grown from glycerol stock in Luria-Bertani nutrient media with carbenicillin (100 μg/mL) for 16 h overnight at 37 °C with shaking. Expression cultures were started by pre-warming Terrific-Broth media with 5% glycerol and carbenicillin (100 μg/mL) to 37 °C before 1% v/v inoculation with the starter culture, then grown for 3 h at 37 °C until addition of IPTG (BioShop, #IPT002) to 0.4 mM for induction. The expression cultures were then transferred to 16 °C and grown for 16 h overnight with shaking, then pelleted with centrifugation and transferred to conical vials for one freeze-thaw cycle at −20 °C. Frozen cell pellets were thawed and resuspended in binding buffer (50 mM HEPES, 500 mM NaCl, 5 mM imidazole, pH 7.2) to a final volume of 100 mL, followed by addition of 0.25 g lysozyme (BioShop, #LYS702). Cell pellet mixtures were sonicated for 25 min (Q700 Sonicator, Qsonica) and clarified by centrifugation to produce a cell lysate.

The REVOLVER was equipped with a 30-mL gravity column containing cobalt-NTA affinity resin and pre-washed with the same binding buffer used to re-suspend the cells for sonication. In parallel, we applied the cell lysates to the pre-auto-homed REVOLVER’s gravity column and let it perform the command sequence <C><P,25,1>< C><P,25,1><C><P,25,1><C><P,11,2><F,5>. The exact protocol was performed by hand in parallel: wash with three 25 mL aliquots of wash buffer, add 11 mL of elution buffer, and collect five 2-mL fractions (the extra 1 mL of elution buffer is added since some of the solution stays in the column that we wanted to account for). The REVOLVER encountered no issues and completed the protocol in 43 minutes.

We then ran an SDS-PAGE analysis of the protein samples and found nearly identical results in terms of the purity of the protein (Fig. 15). The purity is not >95%, which we expect since this protein is difficult to purify and regularly follow-up with anion exchange chromatography or gel filtration. Given to a well-trained biochemist, it would be difficult to discern which of the two panels in Fig. 15 were performed by human and machine.

## 8. Conclusion and future direction

We have designed, built, and characterized open-source systems for parallel protein washing and elution with gravity-column workflows. REVOLVER represents a low cost and highly customizable alternative to more expensive commercial systems, and due to its modular nature can be built in several steps without committing to a full build. This means that the user could build a basic version that does not include a PCB, pumps, or Hall effect sensors, which works essentially as a fraction collector where buffers are added manually. The user can then slowly upgrade the REVOLVER to a point where they have a full MULTI-VOLVER with pumps and Hall sensors. We note that the original design files for all parts of our systems reported here are available to users for ongoing improvements. For example, waste collector version 2 is an example of how to continuously improve our systems, which we recommend users consider over version 1 to improve drainage. Moreover, REVOLVER is programmable via intuitive commands, which makes it highly flexible and reproducible with minimal effort.

There are some additional features that we did not include, but believe would enhance the systems. First, our PCB design has enough extra pins for added functionality, such as: LED status indicators, buzzers, and additional connections to external motor drivers for more pumps for adding cell lysate samples automatically. Second, we chose Arduino Nano microcontrollers for controlling each device because of their low cost and ease of use for users that are already familiar with programming Arduino-like boards. However, using other boards that have internet connectivity, such as the ESP32, would allow for wireless and remote control of each device from a dedicated web server. Third, using a properly designed dedicated user interface (GUI) could make REVOLVER more appealing for the wider community. Finally, while we have made an effort to optimize the design of our parts, different 3D printers with slightly different tolerances might result in pieces that fit loosely after assembly. Nonetheless, we expect the open-source community to contribute and help improve REVOLVER to make it an even better tool.

## Supporting information

Supplementary Table 1 - Bill of materials and tools

## CRediT author statement

**Patrick Diep:** Conceptualization, Methodology, Hardware, Software, Validation, Visualization, Writing – original draft preparation.

**Jose L. Cadavid**: Conceptualization, Methodology, Hardware, Software, Validation, Visualization, Writing – original draft preparation.

**Alexander Yakunin**: Supervision, Writing – reviewing and editing.

**Radhakrishnan Mahadevan:** Supervision, Writing – reviewing and editing.

**Alison P. McGuigan**: Writing – reviewing and editing.

PD and JLC contributed equally to this work and the order of authorship for the manuscript was determined by running the following R code once: if(runif(1) > 0.5){first = “PD”} else {first = “JLC”}. These two authors are free to write their name first in their CVs.

## Acknowledgments

This work was supported by the Ontario Ministry of Economic Development, Job Creation and Trade through the Elements of Bio-mining (EBM) ORF-RE program; as well as the Biochemicals from Cellulosic Biomass (BioCeB) ORF-RE. PD and JLC are recipients of Ontario Graduate Scholarships.

