## Supplementary Figures for "REVOLVER: a low-cost automated protein purifier based on parallel preparative gravity column workflows"

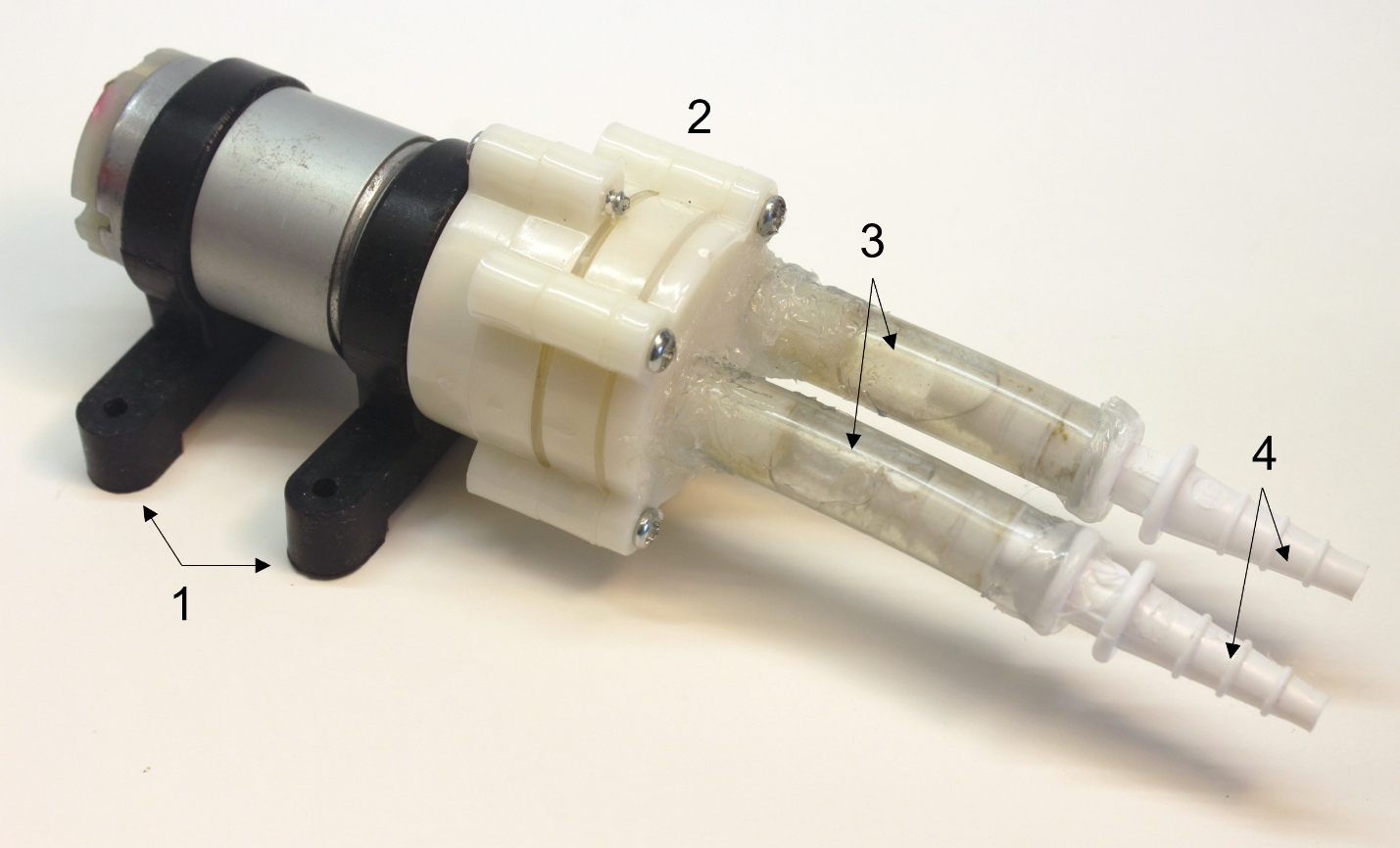

**Supplementary Figure 1. Composite Part – DC Pump with Tube Adaptors.** Assembly Instructions: Step a) attach the plastic sleeves (1) to the DC Pump (2) so it does not roll. Step b) cut two 5 cm segments of large tubing (3) and use silicone to create a water-proof between the DC pump and the large tubing. Step c) use silicone again to attach the tube connectors (4). Allow 24-48 hrs for the silicone to set. Alternatively, one could use hot-glue to connect the pieces, but we found there to be a higher chance for air to be sucked into the tubing through crevices.

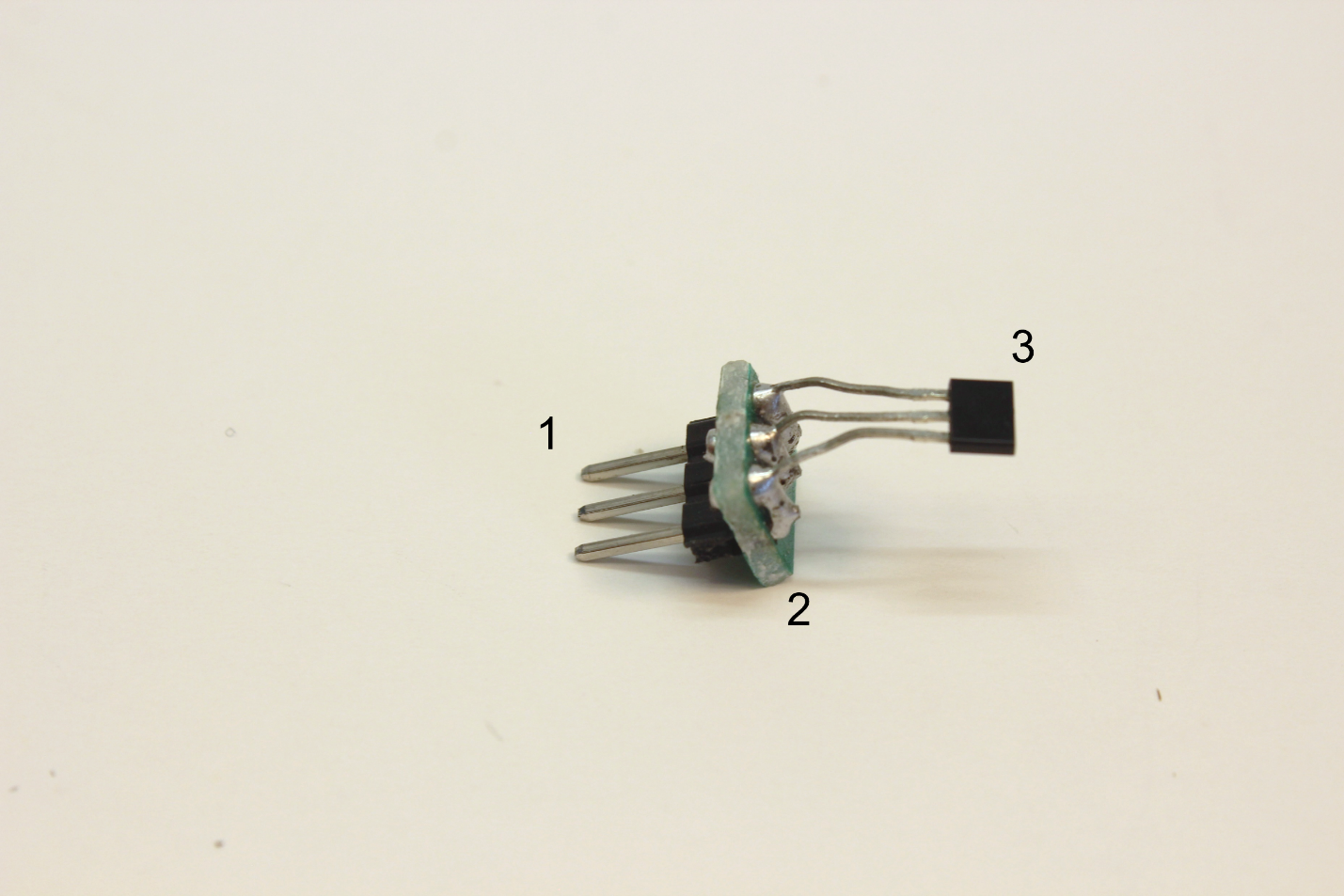

**Supplementary Figure 2. Composite Part – Hall Effect Sensor Board.** Assembly Instructions: Step a) cut a prototyping board (2) to have 2x3 pin holes available. Insert the pin legs of the Hall sensor (3) into one row of the pin holes, then Step b) carefully solder each pin leg to each hole in the row. Step c) insert the short side of a male header pin set (1) into the other row of pin holes, and step d) carefully solder each male pin in the the header to each hole in the row.

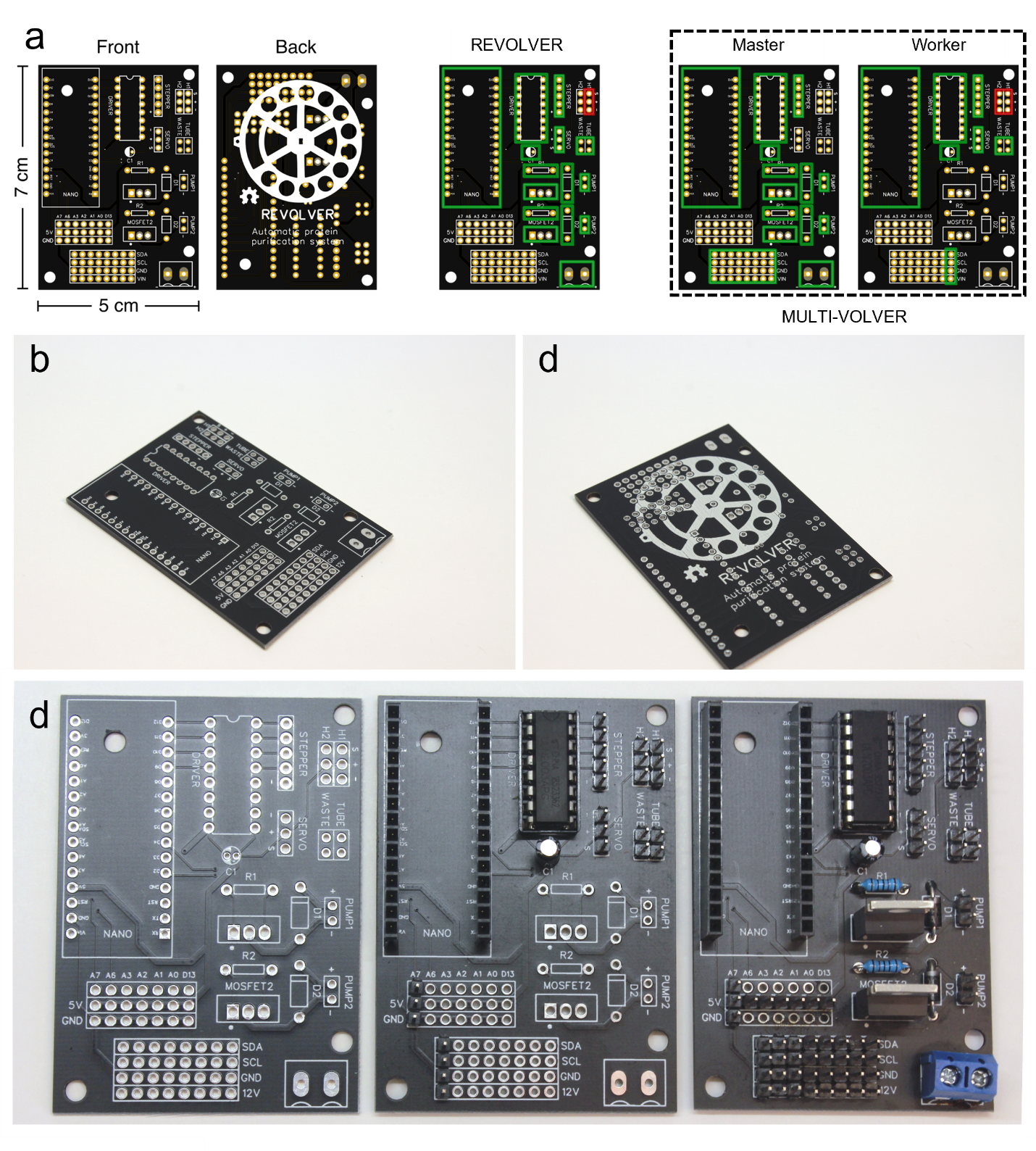

**Supplementary Figure 3. Recommended Print Orientation for REVOLVER Bodies.** (a) Computer-generated visualization of the printed-circuit board (PCB) on EasyEDA. Green boxes indicate the parts of the PCB that will be used in the particular configuration. This includes the Arduino Nano microcontroller, electrical parts that need to be soldered on, and pins connected to the electronics and sensors. Red boxes highlight the need for the REVOLVER to have H1 connected to the waste collectors Hall sensor, and the MULTI-VOLVER to have an additional Hall sensor on the column connected to H2. (b) Front and (c) back of PCB manufactured by JLCPCB. (d) Close-up of (left-panel) empty front, (centre-panel) Worker configuration, (right-panel) Distributor *or* REVOLVER configuration.

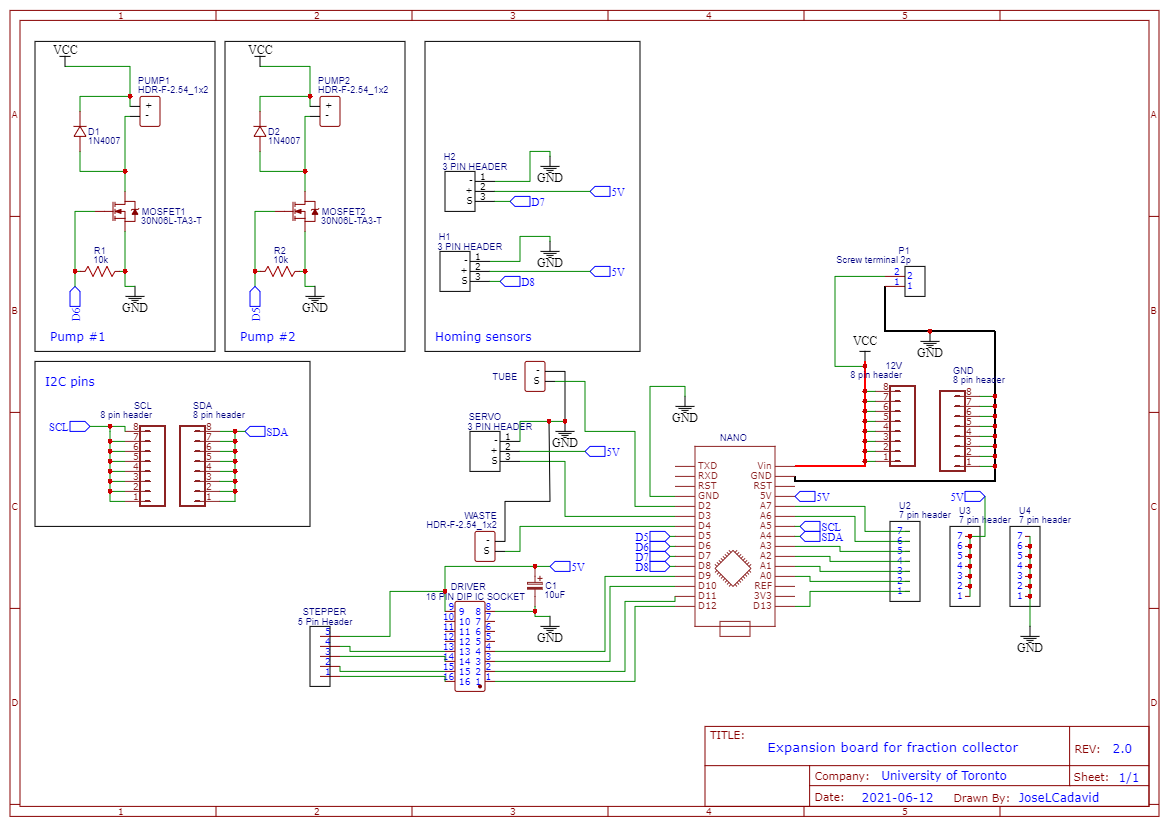

**Supplementary Figure 4. PCB Circuit Diagram from EasyEDA.** This also serves as wiring diagram if the user wishes to use a breadboard prototype first to test/ modify our systems various functionalities.

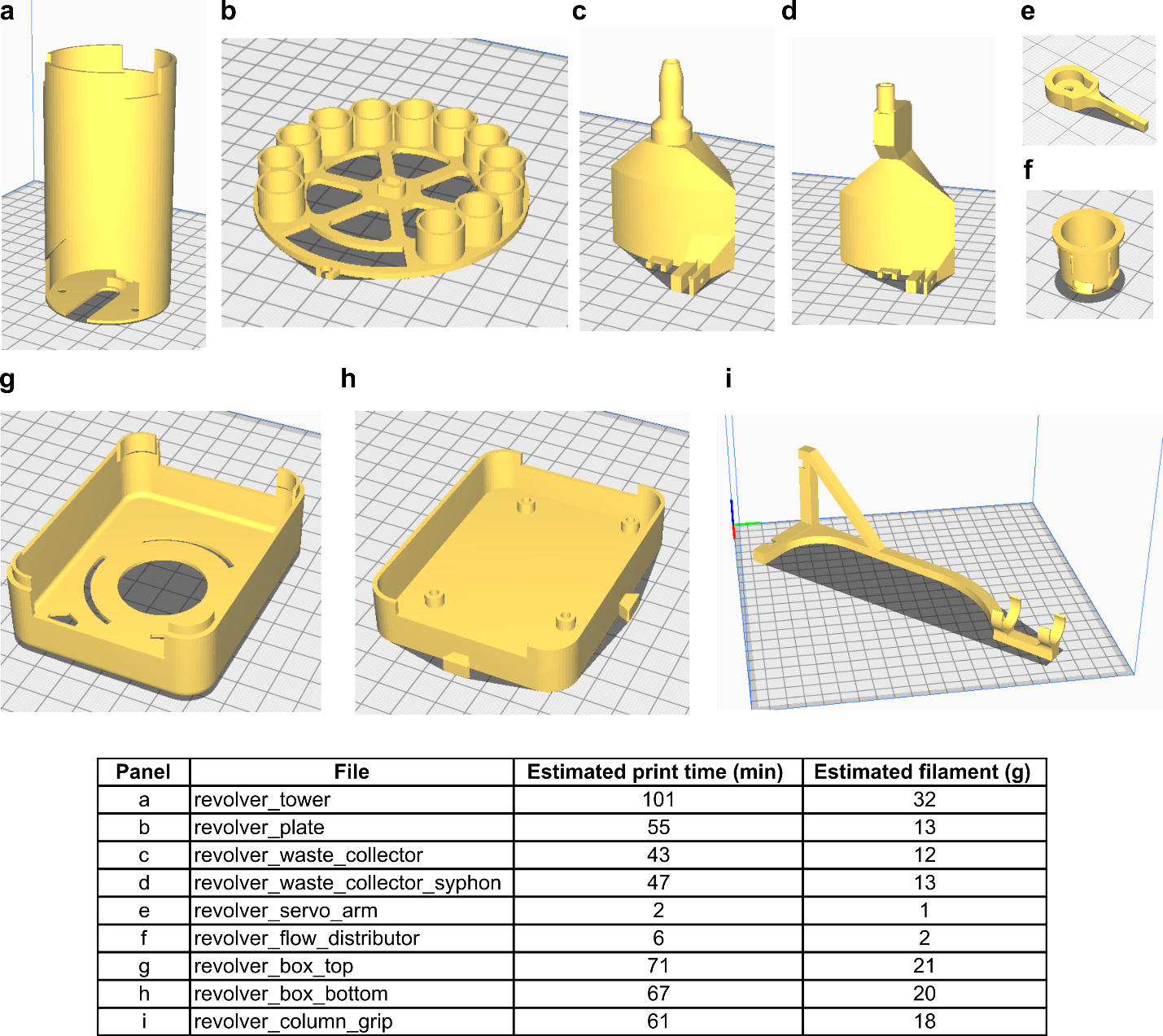

**Supplementary Figure 5. Recommended Print Orientation for REVOLVER Bodies.** These images were generated on Ultimaker Cura’s slicing software. Parts were printer on the Ender 3.

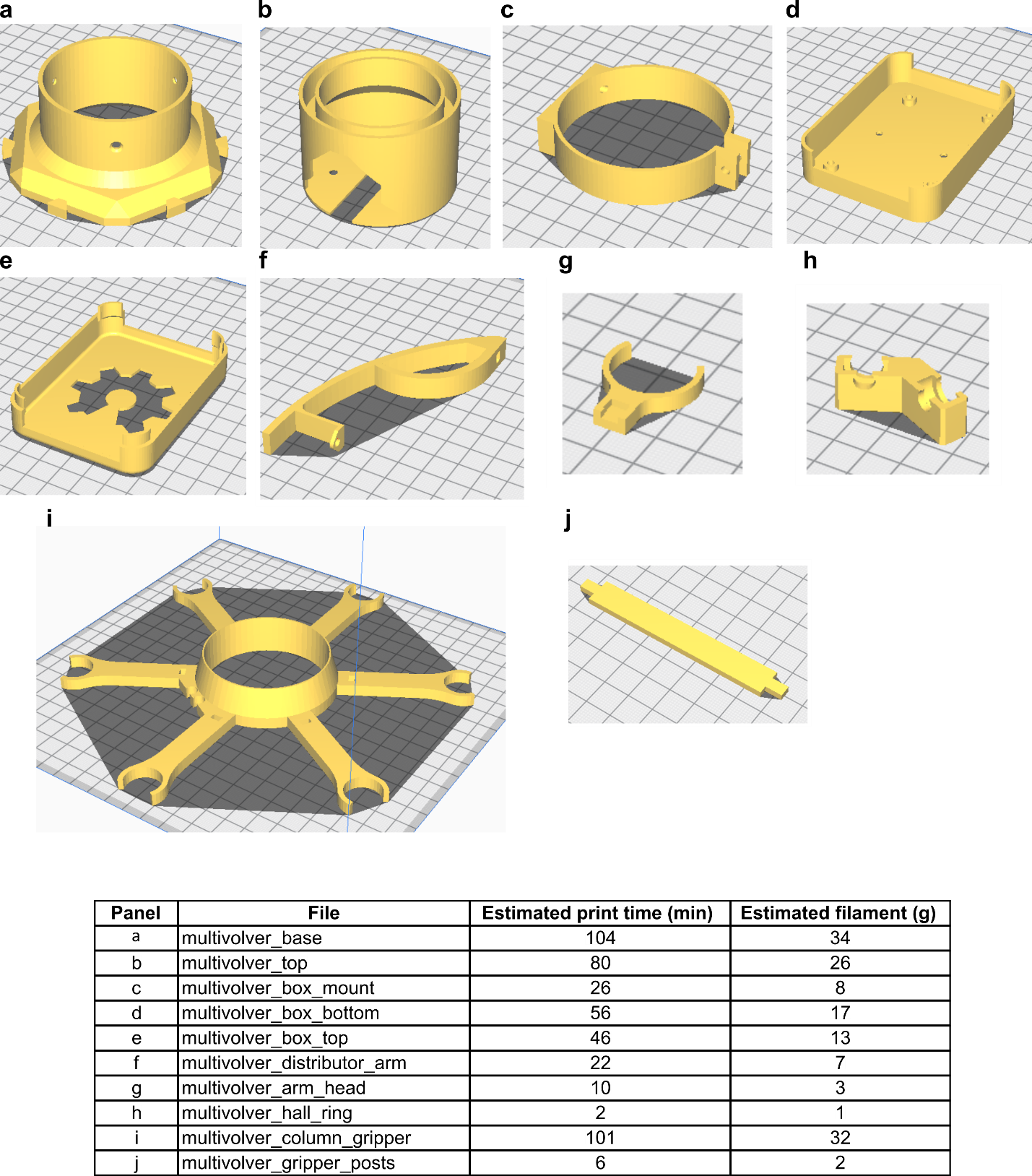

**Supplementary Figure 6. Recommended Print Orientation for MULTI-VOLVER Bodies.** These images were generated on Ultimaker Cura’s slicing software. Parts were printed on the Ender 3.
